# Hub connectivity, neuronal diversity, and gene expression in the *C. elegans* connectome

**DOI:** 10.1101/207134

**Authors:** Aurina Arnatkevičiūte, Ben D. Fulcher, Roger Pocock, Alex Fornito

## Abstract

Studies of nervous system connectivity, in a wide variety of species and at different scales of resolution, have identified several highly conserved motifs of network organization. One such motif is a heterogeneous distribution of connectivity across neural elements, such that some elements act as highly connected and functionally important network hubs. These brain network hubs are also densely interconnected, forming a so-called rich-club. Recent work in mouse has identified a distinctive transcriptional signature of neural hubs, characterized by tightly coupled expression of oxidative metabolism genes, with similar genes characterizing macroscale inter-modular hub regions of the human cortex. Here, we sought to determine whether hubs of the neuronal *C. elegans* connectome also show tightly coupled gene expression. Using open data on the chemical and electrical connectivity of 279 *C. elegans* neurons, and binary gene expression data for each neuron across 948 genes, we computed a correlated gene expression score for each pair of neurons, providing a measure of their gene expression similarity. We demonstrate that connections between hub neurons are the most similar in their gene expression while connections between nonhubs are the least similar. Genes with the greatest contribution to this effect are involved in glutamatergic and cholinergic signalling, and other communication processes. We further show that coupled expression between hub neurons cannot be explained by their neuronal subtype (i.e., sensory, motor, or interneuron), separation distance, chemically secreted neurotransmitter, birth time, pairwise lineage distance, or their topological module affiliation. Instead, this coupling is intrinsically linked to the identity of most hubs as command interneurons, a specific class of interneurons that regulates locomotion. Our results suggest that neural hubs may possess a distinctive transcriptional signature, preserved across scales and species, that is related to the involvement of hubs in regulating the higher-order behaviors of a given organism.

**Author summary:** Some elements of neural systems possess many more connections than others, marking them as network hubs. These hubs are often densely interconnected with each other, forming a so-called rich-club that is thought to support integrated function. Recent work in the mouse suggests that connected pairs of hubs show higher levels of transcriptional coupling than other pairs of brain regions. Here, we show that hub neurons of the nematode *C. elegans* also show tightly coupled gene expression and that this effect cannot be explained by the spatial proximity or anatomical location of hub neurons, their chemical composition, birth time, neuronal lineage or topological module affiliation. Instead, we find that elevated coexpression is driven by the identity of most hubs of the *C. elegans* connectome as command interneurons, a specific functional class of neurons that regulate locomotion. These findings suggest that coupled gene expression is a highly conserved genomic signature of neural hubs that may be related to the specific functional role that hubs play in broader network function.

## Introduction

Neuronal connectivity provides the substrate for integrated brain function. Recent years have seen several systematic and large-scale attempts to generate comprehensive wiring diagrams, or connectomes, of nervous systems [1] in species as diverse as the nematode *Caenorhabditis elegans* [2, 3], *Drosophila* [4, 5], zebrafish [6, 7], mouse [8, 9], rat [10], cat [11], macaque [12, 13], and human [14, 15]. One of the most striking findings to emerge from analyses of these diverse data, acquired using different measurement techniques and at resolution scales ranging from nm to mm, is of a strong conservation of certain topological properties of network organization (reviewed in [1, 16, 17, 18, 19], see also [20]). These properties include a modular organization, such that subsets of functionally related (and usually spatially adjacent) elements are densely interconnected with each other; a hierarchy of modules, such that modules contain nested sub-modules and so on over multiple scales [21, 22]; economical connectivity, such that wiring costs (typically measured in terms of wiring length) are near-minimal given the topological complexity of the system [22, 23]; a heterogeneous distribution of connections across network nodes, such that most nodes possess relatively few connections and a small proportion of nodes have a very high degree of connectivity [3, 24]; and stronger interconnectivity between hub nodes than expected by chance, leading to a so-called rich-club organization (i.e., the hubs are rich because they are highly connected and form a club because they are densely interconnected) [5, 25, 26, 27, 28].

The rich-club organization of hub connectivity is thought to play a central role in integrating functionally diverse and anatomically disparate neuronal systems [26, 29, 30, 31, 32]. Consistent with this view, experimental data and computational modeling indicates that hub nodes, and the connections between them, are topologically positioned to mediate a high volume of signal traffic [33, 34, 35, 36]. This integrative role comes at a cost, with connections between hubs extending over longer distances, on average, than other types of connections, a finding reported in the human [33], macaque [34], rat [37], mouse [38], and nematode [28]. Human positron emission tomography also suggests that hub nodes consume greater metabolic resources and have higher levels of blood flow than other areas [39, 40, 41]. This high metabolic cost may underlie the involvement of hub regions in a broad array of human diseases [17, 29, 31]. Recent work in mice and humans suggests that this high cost of hub connectivity is associated with a distinct gene transcriptional signature. Fulcher and Fornito [38] combined viral tract tracing data on connectivity between 213 regions of the right hemisphere of the mouse brain [8] with *in situ* hybridization measures of the expression of 17 642 genes in each of those regions. They found that transcriptional coupling, across all genes, is stronger for connected compared to unconnected brain regions and that, in general, this coupling decays with increasing separation distance between brain regions. Countering this general trend however, connected pairs of hubs (i.e., the “rich club” of the brain) show the highest levels of transcriptional coupling (compared to hub-nonhub and nonhub pairs), despite being separated by larger distances, on average, and being distributed across diverse anatomical brain divisions. Moreover, this coupling is driven predominantly by genes regulating the oxidative synthesis and metabolism of adenosine triphosphate (ATP), the primary energetic currency of neuronal signaling [42, 43]. Vértes et al. [44] later combined gene expression data of 20 737 genes through 285 cortical areas of the human brain and found evidence that inter-modular hubs in resting state fMRI connectivity networks also have local transcriptional profiles enriched in oxidative metabolism and mitochondria.

Together, the analyses of Fulcher and Fornito [38] and Vértes et al. [44], which were performed using measures of mesoscale (*μ*m to mm) and macroscale (mm to cm) connectivity, respectively, suggest that the molecular signature of hub connectivity, characterized by elevated coupling of genes regulating oxidative metabolism, may be conserved across species and resolution scales. Here, we sought to test this hypothesis by examining the link between gene expression and microscale connectivity in the nematode *C. elegans*. *C. elegans* is the only species to have its connectome mapped almost completely at the level of individual neurons and synapses using electron microscopy [2, 3]. It comprises 302 neurons and around 5600 chemical and electrical synapses [2]. We combined these data on neuronal connectivity with curated information on the binary expression patterns of 948 genes across neurons to examine the relationship between gene expression and the large-scale topological organization of the nematode nervous system. We also used detailed information on neuron spatial positions, birth times, neuronal lineage as well as the functional and chemical composition of each neuron to understand the mechanisms through which gene expression might influence network topology. Paralleling findings in humans and mouse, we find that hub neurons of *C. elegans* are genomically distinct, with connected hub neurons showing the most similar patterns of gene expression. Genes that contribute most to this effect are involved in regulating glutamate and acetylcholine function, and neuronal communication. We demonstrate that this effect is not explained by factors such as neuronal birth time, lineage, neurotransmitter system or spatial position but is rather related to the functional specialization of hub neurons in mediating higher order behaviours of the organism.

## Materials and methods

We first describe the neural connectivity data used to construct a connectome for *C. elegans* and the methods used to quantify network connectivity and other properties of individual neurons, including their neurochemical composition, birth times, and lineage relationships. We then describe the gene expression data, how we measure expression similarity between pairs of neurons, and our method for scoring the contribution of individual genes to patterns of correlated gene expression. Note that all data used for analysis in this work were obtained from publicly available sources, and can be downloaded from an accompanying figshare repository (**figshare.com/s/797199619fbabdab8c86**). Code to process this data and reproduce all figures and analyses presented here is on github (**github.com/BMHLab/CElegansConnectomeGeneExpression**).

### Neuronal connectivity data

The nervous system of *C. elegans* comprises 302 neurons, divided into the pharyngeal nervous system (20 neurons) and the somatic nervous system (282 neurons). The spatial positions of neurons, and their interconnections, are genetically determined and highly reproducible across organisms [45]. Neuronal connectivity data for the adult hermaphrodite *C. elegans* was first mapped by White et al. [2] through detailed electron microscopy, and then revised by Chen et al. [46] and Varshney et al. [3]. Here we analyze the larger somatic nervous system using data from Varshney et al. [3], who mapped connectivity between 279 neurons (282 somatic neurons, i.e., excluding CANL/R, and VC6, which do not form synapses with other neurons), which we obtained from WormAtlas [47] (**www.wormatlas.org/neuronalwiring.html#NeuronalconnectivityII**).

Connectivity data are available for both electrical gap junctions and chemical synapses. The chemical synapse network is both directed (i.e., the pre-synaptic and post-synaptic neurons are identified) and weighted (as the number of synapses from one neuron to another), while gap junctions are conventionally represented as weighted (as the number of electrical synapses connecting two neurons), undirected connections. Previous investigations of *C. elegans* neuronal connectivity have used differently processed versions of these data, including: (i) only chemical synapses [48]; (ii) a combination of chemical and electrical synapses as a directed network (electrical synapses represented as reciprocal connections) [49, 50]; (iii) a combination of chemical and electrical synapses as an undirected network (representing unidirectional and reciprocal chemical connections equivalently) [28, 51, 52, 53]; or (iv) comparing multiple connectome representations [54]. Our analysis here focuses on the combined directed, binary network, treating gap junctions as bidirectional connections. The resulting connectome contains 279 neurons, with 2 990 unique connections linking 2 287 pairs of neurons. Note that the qualitative results presented here are not highly sensitive to the connectome representation. For example, we observe similar results when analyzing just the directed chemical connectivity network.

In addition, we assembled a range of data characterizing other properties of *C. elegans* neurons. To examine the effect of physical distance between pairs of neurons, we obtained two dimensional spatial coordinates for each neuron as `celegans277.mat` from **www.biological-networks.org/?page_id=25** [55]. Coordinates for three neurons (AIBL, AIYL, SMDVL) were missing in this dataset, and were reconstructed by assigning identical coordinates to the corresponding contralateral neurons (AIBR, AIYR, SMDVR) [56]. To examine the influence of anatomical location, each neuron was labeled by its anatomical location, as: (i) ‘head’, (ii) ‘tail’, or (iii) ‘body’, using data from release WS256 of WormBase [57], (**ftp://ftp.wormbase.org/pub/wormbase/releases/WS256/ONTOLOGY/anatomy_association.WS256.wb**). These annotations were assigned to individual neurons using the anatomical hierarchy defined in WormBase, which we retrieved using the WormBase API (WormMine: **intermine.wormbase.org**) [57], propagating each term down the hierarchy to individual neurons. We manually corrected the assignment of twelve head neurons (ALA, AVFL, AVFR, AVG, RIFL, RIFR, RIGL, RIGR, SABD, SABVL, SABVR, SMDVL), which were assigned as ‘head’ in WormAtlas [47] but not on WormBase. To examine the influence of neuronal subtype, all neurons were labeled to one (or multiple) of the following three categories: (i) ‘sensory’ (support receptive function), (ii) ‘motor’ (contain neuromuscular junctions), or (iii) ‘interneuron’ (all other neurons) [2]. A total of 79 neurons are annotated as sensory, 121 annotated as motor, and 97 annotated as interneurons (including eighteen neurons assigned to two categories: five are both ‘interneuron’ and ‘sensory’, seven are ‘interneuron’ and ‘motor’, and six are both ‘sensory’ and ‘motor’ neurons). To examine the neurotransmitter systems used by each neuron, neurons were labeled by matching to Table 2 of Pereira et al. [58]. In order to determine the influence of neuron birth time, we obtained neuronal birth time information in minutes from the Dynamic Connectome Lab website (**https://www.dynamic-connectome.org/?page_id=25**), [56]. To assess the influence of lineage similarity, we obtained a measure of lineage distance for all pairs of neurons from previously published embryonic and post-embryonic lineage trees [59, 60], using data downloaded from WormAtlas (**http://www.wormatlas.org/neuronalwiring.html#Lineageanalysis**) [47]. In this dataset, the closest common ancestor neuron was identified for each pair of neurons, and then the lineage distance was calculated as the number of cell divisions from the closest common progenitor neuron.

### Network analysis

In this section we describe the network analysis methods used to characterize the *C. elegans* connectome.

#### Degree

Neuronal connectivity is most simply quantified by counting the number of neurons that a given neuron projects to, known as its out-degree, *k*_out_, or by counting the number of neurons that a given neuron receives projections from, known as its in-degree, *k*_in_. These quantities can be summed to give the total number of connections involving a given neuron, *k*_tot_ = *k*_in_ + *k*_out_, which we refer to as simply the degree, *k*, throughout this work. At a given degree threshold, *k*, each neuron was classified as either a ‘hub’ (degree > *k*) or a ‘nonhub’ (degree ≤ *k*). All connections were subsequently classified in terms of their source and target neurons as either ‘rich’ (hub → hub, or hub ↔ hub), ‘feed-in’ (nonhub → hub), ‘feed-out’ (hub → nonhub), or ‘peripheral’ (nonhub → nonhub, or nonhub ↔ nonhub).

#### Rich-club organization

We used the rich-club coefficient, *ϕ*(*k*), to quantify the interconnectivity of hub neurons:

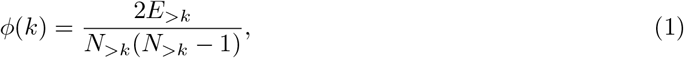

where *N*_>*k*_ is the number of nodes with degree > *k*, and *E*_>*k*_ is the number of edges between them [61]. Thus, *ϕ*(*k*) measures the link density in the subgraph containing nodes with degree > *k*. Because *ϕ*(*k*) invariably increases with *k* (as nodes with higher degree make more connections, yielding a higher expected link density in the subgraph containing nodes with degree > *k*), we compared *ϕ*(*k*) measured from the *C. elegans* connectome to the mean value of an ensemble of randomized null networks, *ϕ*_rand_(*k*) [61]. An ensemble of 1000 null networks was generated by shuffling the links in the empirical network while retaining the same degree sequence [62] (rewiring each edge an average of 50 times per null network) using the `randmio_dir` function from the Brain Connectivity Toolbox [63]. The normalized rich-club coefficient, Φ_norm_(*k*), was computed as the ratio of the rich-club coefficient of the empirical network to the mean rich-club coefficient of the ensemble of randomized networks:

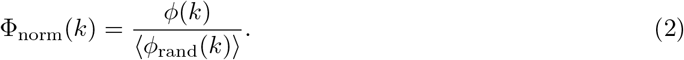

Values of Φ_norm_ > 1 indicate rich-club organization of the network, with statistically significant deviations assessed by computing a *p*-value directly from the empirical null distribution, *ϕ*_rand_(*k*), as a permutation test [25].

#### Modularity

To investigate whether our results were influenced by the well-known modular organization of the *C. elegans* connectome [22, 51, 52, 54, 64], we decomposed the network into modules using two methods. The first method involved applying the Louvain community detection algorithm [65] using the `community_louvain` function in the Brain Connectivity Toolbox [63]. To identify the optimal modular assignment, neurons were assigned to modules using consensus clustering across 1000 repeats of the Louvain algorithm [66], weighting each partition by its modularity, *Q*, using the `agreement_weighted` and `consensus_und` functions of the BCT[63]. The second modular decomposition was taken from a previously-reported nine-module partition derived from an Erdös-Rényi Mixture Model (ERMM) [52].

### Gene expression

Gene expression is represented as a binary indicator of which genes are expressed in a given neuron using data from many individual experiments compiled into a unified data resource on WormBase [57]. We use release WS256 of this dataset (**ftp://ftp.wormbase.org/pub/wormbase/releases/WS256/ONTOLOGY/anatomy_association.WS256.wb**) and analyze annotations made ‘directly’ to individual neurons, excluding ‘uncertain’ annotations (see Supplementary Information for details on annotations). We denote the expression of a gene in a neuron as a ‘1’, and other cases as a ‘0’. Note that a value of ‘0’ indicates either: (i) ‘gene is not expressed’ or (ii) ‘there is no information on whether gene is expressed’. Expression data are sparse, in part due to incomplete annotations—an average of 30 genes are expressed in each neuron (range: 3 to 138 genes), and each gene is expressed in an average of 9 neurons (range: 1 to 148 neurons). A total of 948 genes are expressed in at least one neuron, allowing us to represent the full expression dataset as a binary 279 (neuron) × 948 (genes) matrix, shown in Fig. 1C.

**Fig 1.**
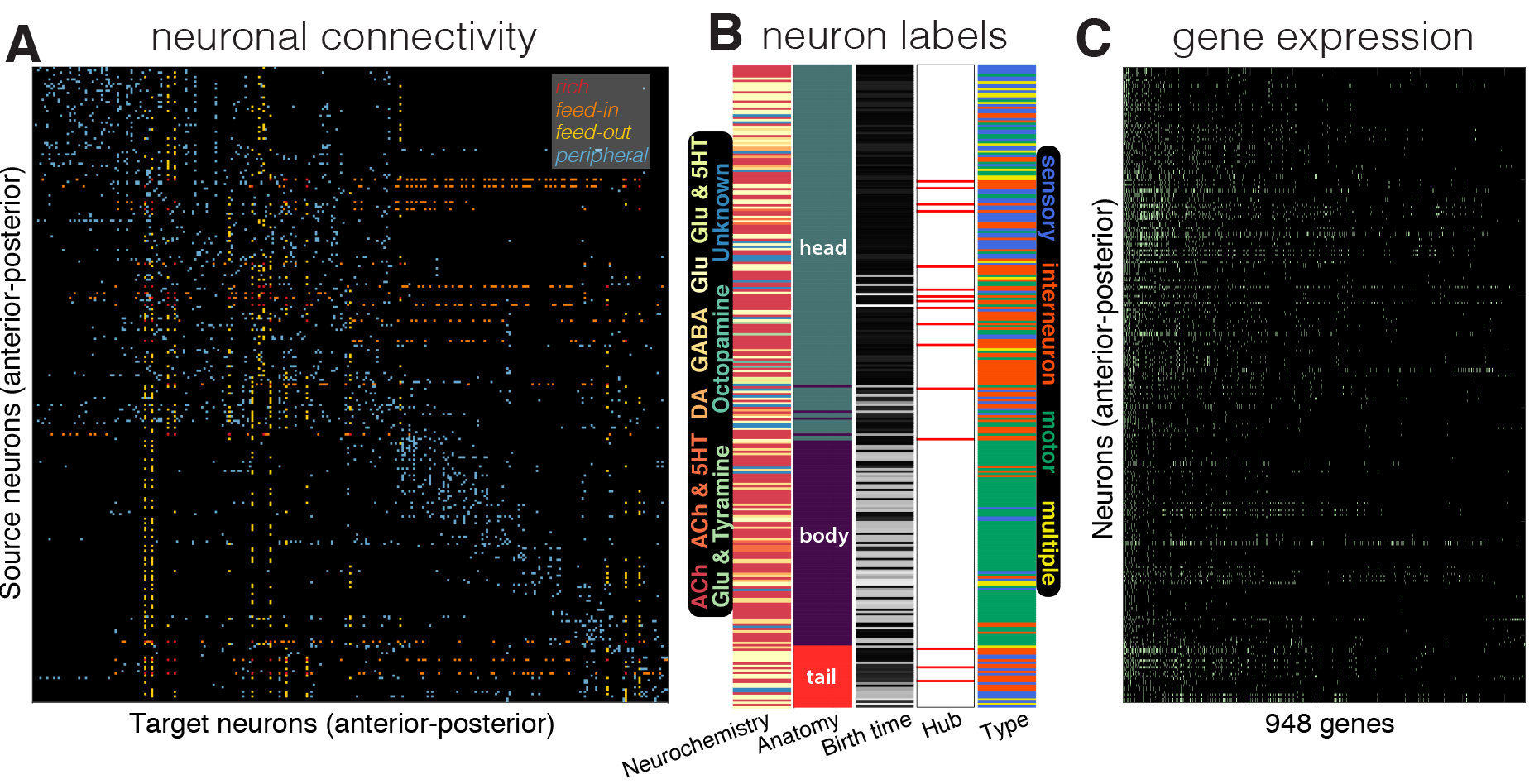
Schematic representation of the data used in this study. All plots show neurons (rows) ordered by anterior-posterior along the longitudinal axis, from the top of the head (upper, left) to the bottom of the tail (lower, right). (A) Connectivity matrix summarized 2990 directed chemical and electrical connections between 279 neurons from neuron *i* (row) to neuron *j* (column). Connections are colored according to how they connect hubs (*k* > 44) and nonhubs (*k* ≤ 44), as ‘rich’ (hub → hub), ‘feed-in’ (nonhub → hub), ‘feed-out’ (hub → nonhub), and ‘peripheral’ (nonhub → nonhub). (B) Neurochemistry (types as labeled), anatomical location (as labeled), birth time (from early born neurons, black, to late-born neurons, white), hub assignment (hubs labeled red), and functional type (as labeled). (C) Binary gene expression indicated as a green dot when a gene (column) is expressed in a neuron (row).

### Correlated gene expression

Our primary aim in this work is to understand how pairwise patterns of neuronal connectivity (shown in Fig. 1A) relate to coupled expression across 948 genes between pairs of neurons (i.e., pairs of rows of Fig. 1C). To estimate the coupling between neuronal gene expression profiles, we required a similarity measure for pairs of neurons that captures their similarity of gene expression profiles, or correlated gene expression (CGE). We used a binary analogue of the linear Pearson correlation coefficient, the mean square contingency coefficient [67]:

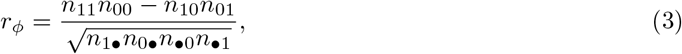

for two vectors, *x* and *y*, of length *L*(= 948), where *n_xy_* counts the number of observations of each of the four outcomes (e.g., 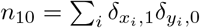 counts the number of times *x* = 1 and *y* = 0), while the symbol • sums across a given variable (e.g., 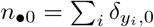 counts the number of times *y* = 0). The coefficient assumes a maximum value *r_ϕ_* = 1 when *x* and *y* are identical (such that *n*_11_ + *n*_00_ = *L*), and a minimum value *r_ϕ_* = −1 when *x* and *y* are always mismatched (such that *n*_10_ + *n*_01_ = *L*).

One concern about applying this measure to sparsely annotated data is that it may be biased by differences in the number of expressed genes in a neuron, which ranged from 3 (0.3% of 948 genes analyzed here) to 138 (14.6%). To explore this further, we compared *r_ϕ_* with several other commonly used metrics of association between binary vectors including Yule’s *Q* and Jaccard. While these other binary correlation metrics exhibited strong bias with the proportion of gene expression annotations made to a given neuron, *r*_*ϕ*_ was not biased (see Fig. S1). Note that we use *r*_*ϕ*_ to denote this coefficient given that it is commonly referred to as the phi coefficient in the literature; this notation should not be confused with *ϕ*(*k*), which is used to denote the rich club coefficient.

The 92 bilateral pairs of neurons (e.g., AVAL/AVAR, CEPVL/CEPVR, etc.) exhibit highly correlated gene expression patterns, with all bilateral pairs of neurons exhibiting *r*_*ϕ*_ > 0.8, and 96% of bilateral pairs exhibit *r_ϕ_* > 0.95. Although including bilateral pairs of neurons do not change the main results of this paper, to ensure that our results are not driven by the high CGE between bilateral pairs of neurons, we excluded CGE values of bilateral pairs of neurons from all analyses involving CGE.

### Gene scoring and enrichment

Our CGE measure, *r_ϕ_*, quantifies the association between the expression profiles of two neurons across all 948 genes. To further investigate the role of individual genes in producing different CGE patterns, we developed a method for scoring the contribution of each individual gene to the overall correlation coefficient, following prior work using continuous gene expression data [38]. Performing similar analyses with *C. elegans* data poses additional challenges due to: (i) binary expression data, making robust quantification difficult; (ii) sparse and incomplete data, posing statistical problems for quantifying a signal in genes with limited expression; and (iii) low genome coverage (less than 5% of the protein coding genes in *C. elegans*), constraining our ability to perform a comprehensive enrichment analysis, e.g., using the Gene Ontology (GO) [68].

Given that *r*_*ϕ*_ [Eq. 3] treats mutual gene expression (i.e., cases in which a gene is jointly expressed in both neurons) (*n*_11_) the same as mutual absence of gene expression (*n*_00_), we started by developing a new analytic measure of the probability of mutual gene expression, given its clearer biological interpretation (see Supporting Information). This measure was not biased by differences in the relative number of expressed genes (Fig. S1D) and yields qualitatively similar outputs to *r*_*ϕ*_ on our data. Thus, while we use *r*_*ϕ*_ throughout this work for its ease of interpretation (as an analogue of the conventional correlation coefficient), our new probability-based CGE measure allowed us to motivate a method for quantifying the contribution of individual genes (and functional groups of genes) to patterns of CGE that addresses some of the above-mentioned challenges. Note that our basic findings replicated regardless of whether we use *r*_*ϕ*_ or this new measure to quantify CGE (e.g., cf. Fig. S2).

As a starting point, we quantified the contribution of individual genes to differences in CGE for different categories of neuron pairs, specifically for (i) increased CGE in connected compared to unconnected pairs of neurons, and (ii) increased CGE in rich and feeder compared to peripheral connections. We first scored each gene for whether it is more likely to be expressed in a given class of neuron pair over another as the probability of obtaining at least as many matches (defined as expression in both neurons of a pair) as observed under a random CGE null model using the binomial distribution:

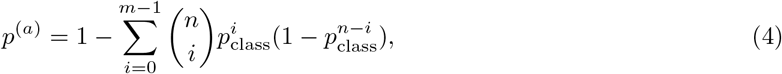

where *m* is the number of matches (a match indicates that a given gene was expressed in both neurons) on the class of neuron pairs of interest, *n* is the total number of matches across all neuron pairs considered in the analysis, *p*_class_ = *n*_class_/*M* is the probability of the given class of inter-region pairs, as the total number of neuron pairs of that class, *n*_class_, divided by the maximum number of possible neuron pairs, *M*, for a given gene, indexed with *a*. This score, *p*^(*a*)^, can be interpreted as a *p*-value under the null hypothesis that the number of expression matches of gene *a* is consistent with a purely random pattern of matches/mismatches across edges, giving lower values to genes with more matches in the edge class of interest (compared to an alternative set of edges) than expected by chance. For reasons described earlier, bilateral pairs of neurons were excluded from all scoring procedures and, to ensure that each gene contributes a meaningful score, we imposed a data quality threshold on the number of possible matches, *n* ≥ 10.

Our first analysis compares two mutually exclusive types of neuron pairs: (i) all pairs of neurons that are connected by at least one chemical or electrical synapse, and (ii) all pairs of neurons that are unconnected. For this analysis, *p*_class_ = 0.059 is the proportion of neuron pairs that are connected, *n* is the total number of neuron pairs that both exhibit expression of gene *a*, and *m* is the number of neuron pairs that are structurally connected for which both neurons express gene *a*. A total of 414 (/948) genes had *n* ≥ 10 for this analysis. Our second analysis compares pairs of connected neurons for which at least one is a hub (i.e., rich, feed-in, or feed-out connections), to pairs in which both neurons are nonhubs (i.e., peripheral connections). In this case, *p*_class_ = 0.35 is the proportion of connected pairs of neurons that involve hubs, *n* is the number of connected neuron pairs for which gene *a* is expressed in both, and *m* counts the number of connected neuron pairs involving hubs for which gene *a* is expressed. A total of 168 (/948) genes had sufficient data, *n* ≥ 10, for this analysis.

As well as interpreting the list of individual genes that were significant after correcting for multiple hypothesis comparison, we performed an over-representation analysis (ORA) using the genes that contribute most to a given connectivity pattern to assess whether any gene sets (GO categories) were statistically over-represented in this list. To obtain the gene list, we used the false discovery rate correction of Benjamini and Hochberg [69] on *p*-values computed using Eq. (4), which were thresholded at a stringent level of *p* = 10^−4^ (corresponding to approximately the top 20% of genes in each analysis). Over-representation for each biological process GO category with 5 to 100 genes available was quantified as an FDR-corrected *p*-value (across around 700 GO categories) using version 3.0.2 of *ErmineJ* software [70]. Biological process GO annotations [68] were obtained from GEMMA [71] (using **GenericwormnoParents.an.txt.gz** downloaded on March 31, 2017). Gene Ontology terms and definitions were obtained in RDF XML file format downloaded from **archive.geneontology.org/latest-termdb/go_daily-termdb.rdf-xml.gz** on March 31 2017.

## Results

Our main aim in this work is to relate patterns of pairwise connectivity in the *C. elegans* neuronal connectome to correlated gene expression, focusing particularly on hub connectivity. A schematic overview of our data is in Fig. 1, including the directed binary connectome (Fig. 1A), additional anatomical data gathered for each neuron (Fig. 1B), and binary gene expression across 948 genes (Fig. 1C).

Our analysis is presented in five parts: (i) given past evidence for a major effect of physical distance on connection probability and CGE [38], we first characterize the spatial dependency of connection probability and CGE; (ii) we confirm the rich-club organization of the *C. elegans* connectome; (iii) we show that CGE is increased in connected pairs of neurons relative to unconnected pairs, in electrical synapses relative to chemical synapses, and in connected hub neurons relative to other types of connected neuron pairs (mirroring previous results in the mesoscale mouse connectome [38]); (iv) we demonstrate that high CGE between connected hub neurons is not driven by factors like stereotypical interneuron expression, birth time, lineage similarity, neuromodulator types or expression similarity within modules, but may be driven by the high CGE of command interneurons; (v) we characterize the contribution of specific genes, and broader gene ontology categories, to the observed patterns.

### Spatial dependency

Previous work has demonstrated the importance of spatial effects in driving patterns of gene expression, with more proximal brain areas exhibiting more similar gene expression patterns than more distant brain areas [38, 72, 73, 74]. Connection probability also decreases with spatial separation between: brain areas [38, 75, 76, 77, 78, 79], individual neurons in mouse primary auditory cortex [80], and neurons of the *C. elegans* nervous system (cf. Fig. S1 of [49]). Unlike network analyses of mammalian brains, where all neurons are confined to a spatially contiguous organ, neurons of the *C. elegans* nervous system are distributed throughout the entire organism, forming a dense cluster of 147 neurons in the head (all within 130 *μ*m), 105 sparser neurons in the body (spanning 1.02 mm), which are predominantly motor neurons (75%), and another dense cluster of 27 neurons in the tail (all within 90 *μ*m of each other), as plotted in Fig. 2. In order to examine the relationship between connectivity and CGE, we first need to understand the spatial dependence of both connectivity and CGE to characterize the extent to which previously reported spatial dependencies of these measurements apply to the microscale nervous system of *C. elegans*.

**Fig 2.**
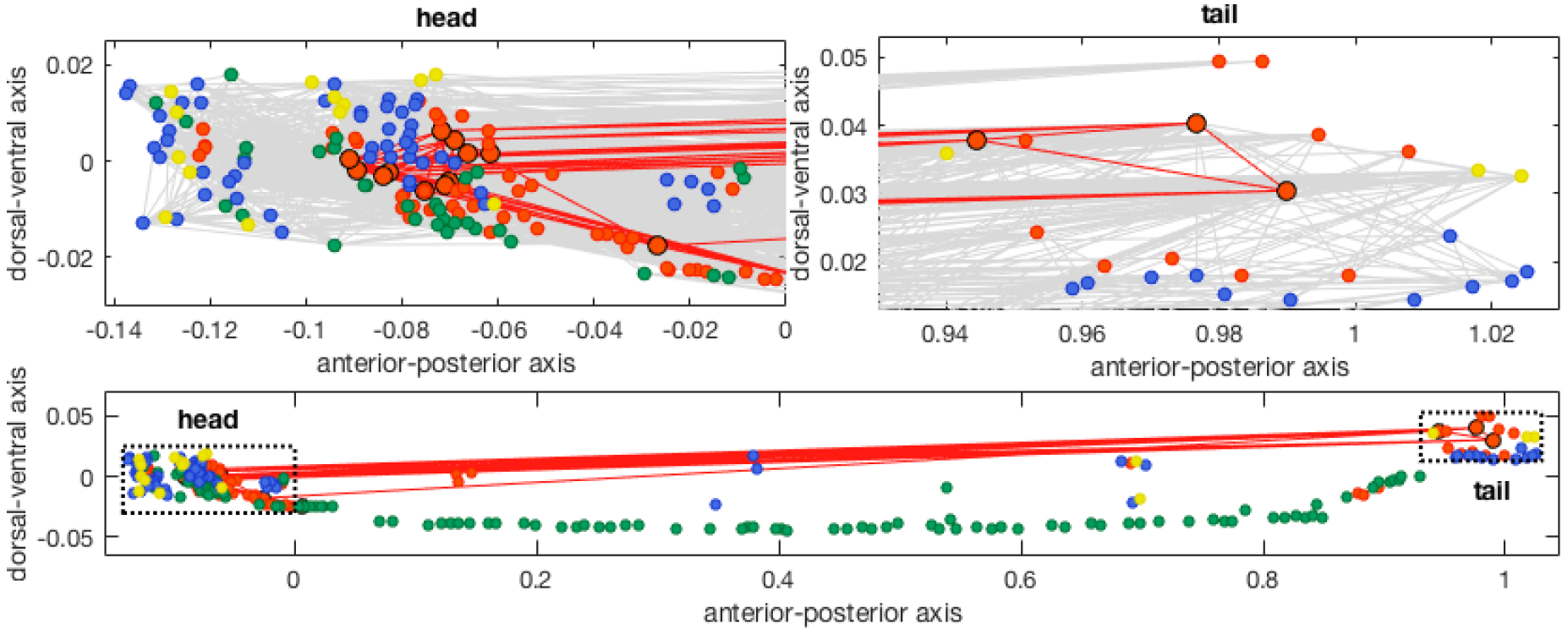
Hub neurons are contained within the head and tail of *C. elegans*. Neurons are positioned along the anterior–posterior (horizontal), and dorsal–ventral (vertical) axes, and are colored by type: (i) interneuron (85 neurons, orange), (ii) sensory (68 neurons, blue), (iii) motor (108 neurons, green), or (iv) multiple assignments (18 neurons, yellow). Hub neurons (i.e., neurons with *k* > 44, see Fig. 5) are shown as larger circles and outlined in black. ‘Rich-club’ connections between hub neurons are shown (red), and all other connections are also shown in the upper plots (gray). Axes of each subplot are to scale with each other, and the upper zoomed-in plots of the head and tail are shown as dotted rectangles in the lower plot.

We first characterize the probability that two neurons will be connected given their source and target types, labeling each neuron as being in either the ‘head’, ‘body’, or ‘tail’ of *C. elegans*. Connection probability is plotted as a function of Euclidean separation distance in Fig. 3 for each combination of source and target neuron labels, across 10 equiprobable distance bins (with exponential fits added for visualization). Distinguishing connections by source and target neuron types uncovers clear spatial relationships (that are obscured when all connections are grouped, as in [49]), that differ across connection classes. From the very short distance scale of ⪅ 100 *μ*m of head → head and tail→tail connections to the very longest-range head→tail and tail→head connections (⪆ 1 mm), connection probability decreases with separation distance (Fig. 3A). For connections between pairs of neurons located in the body, ranging up to ≈ 1 mm, a near-exponential trend is exhibited, mirroring results in other species and across longer length scales [76], including mouse [38, 81], and in rodents and primates [75]. Other connections do not exhibit strong spatial connectivity relationships, i.e., connections between the body and head or between the body and tail, shown in Fig. 3B.

We next investigate the dependence of CGE, *r*_*ϕ*_, on the separation distance between neuron pairs, shown in Fig. 4. CGE decreases slightly with separation distance for the spatially close neurons within the head (Fig. 4A) and within the tail (Fig. 4B), but not for pairs of neurons involving the body (Fig. 4C). The decreasing trend in CGE with distance within the head and tail is primarily driven by a subset of nearby neurons with high *r*_*ϕ*_. It may therefore represent a relationship specific to particular functionally related neurons, rather than a general, bulk spatial relationship seen in macroscopic mammalian brains [38]. Accordingly, attempting to correct for a bulk, non-specific trend by taking residuals from an exponential fitted to the relationship produced artifactual reductions in the CGE of many neuron pairs (shown in Fig. S3). Thus, we found no evidence for bulk spatial relationships of *r*_*ϕ*_ in the neuronal connectome of *C. elegans*.

### Hub connectivity

Next we analyze the topological properties of the *C. elegans* connectome, represented as a directed, binary connectivity matrix between 279 neurons, combining directed chemical synapses and undirected electrical gap junctions (Fig. 1). The degree distribution is shown in Fig. 5A, where neurons are distinguished by type: 68 sensory neurons, 85 interneurons, 108 motor neurons, and 18 neurons with multiple assignments. Consistent with the results of Towlson et al., who used an undirected version of the connectome (by ignoring the directionality of chemical synapses), we found a positively-skewed degree distribution containing an extended tail of high-degree hub interneurons. Hub interneurons of *C. elegans* are mostly command interneurons and mediate behaviors like coordinated locomotion and foraging [82].

Using the normalized rich-club coefficient, Φ_norm_, to quantify the extent to which hubs are densely interconnected, we confirmed the results of Towlson et al. [28], finding rich-club organization in the connectome, as shown in Fig. 5B. The figure plots the variation of Φ_norm_ across degree thresholds, *k*, at which hubs are defined (as neurons with degree > *k*), with red circles indicating a significant increase in link density among hubs relative to 1000 degree-preserving nulls (permutation test, *p* < 0.05). The plot reveals rich-club organization (Φ_norm_ > 1) at the upper tail of the degree distribution, particularly across the range 44 < *k* < 63, shaded gray in Fig. 5B. Similar results were obtained using weighted representations of the connectome (i.e., using information about the number of synapses in the connectivity network) for two different definitions of the weighted rich-club coefficient [83], shown in Fig. S4. Throughout this work, we define a set of hubs as the sixteen neurons with *k* > 44, which corresponds to the lowest degree threshold at which the network displays a contiguous region of significant rich-club organization at high *k*. Our list of hubs includes all of the 11 hub neurons of Towlson et al. [28] at 3*σ* (see Fig. S1) with five additional hubs identified in our analysis of the directed connectome.

The rich-club connectivity of the *C. elegans* connectome is accompanied by an increase in mean hub-hub connection distance [28], with a significant increase through the topological rich-club regime (right-tailed Welch’s *t*-test, *p* < 0.05), shown in Fig. 5B. This can be attributed to a relative increase in long-distance hub-hub connections between the head and tail, shown in Fig. 2 (cf. Fig. S5). The high connection density and long mean anatomical distance between pairs of hub neurons counters the general trend in the *C. elegans* connectome, where the probability of connectivity between two neurons decays with their separation distance (Fig. 3). These results are consistent with previous findings across diverse neural systems and suggest that the rich club may provide a central yet costly backbone for neuronal communication in *C. elegans* [28, 33].

**Fig 3.**
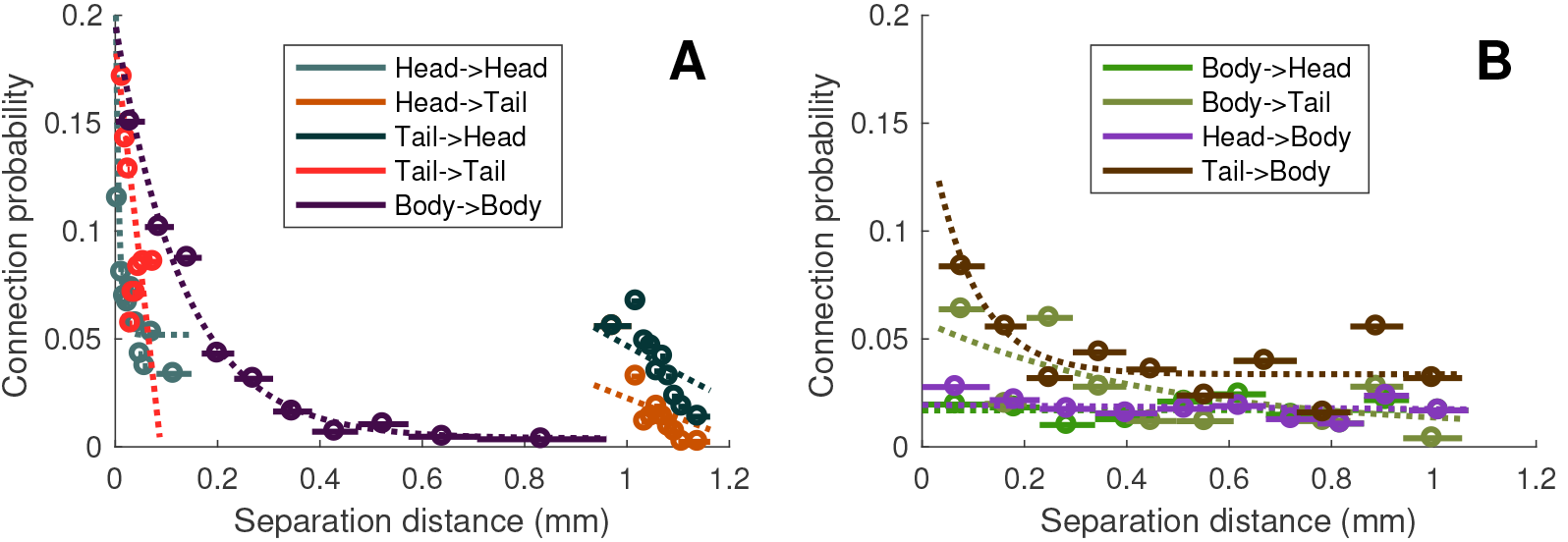
Connection probability decreases with separation distance within and between the head and tail, and within the body. The connection probability for a pair of neurons as a function of their Euclidean distance is estimated in 10 equiprobable distance bins, shown as a circle (bin centers) and a horizontal line (bin extent). There is a decreasing relationship for connections: within the head (aqua), from head→tail (brown) and from tail→head (stone blue), within the tail (red), and within the body (dark purple). Exponential fits of the form *f*(*x*) = *A*exp(−*λx*) + *B*, some of which appear approximately linear across the range of the data, are shown as dotted lines. (B) Plots as in (A), but for connection classes between the body and head/tail: from body→head (forest green), from body→tail (dirt green), connections from head → body (purple), and from tail→body (dark brown). Apart from a small effect at short range for tail→body connections, these connection classes show minimal distance dependence.

**Fig 4.**
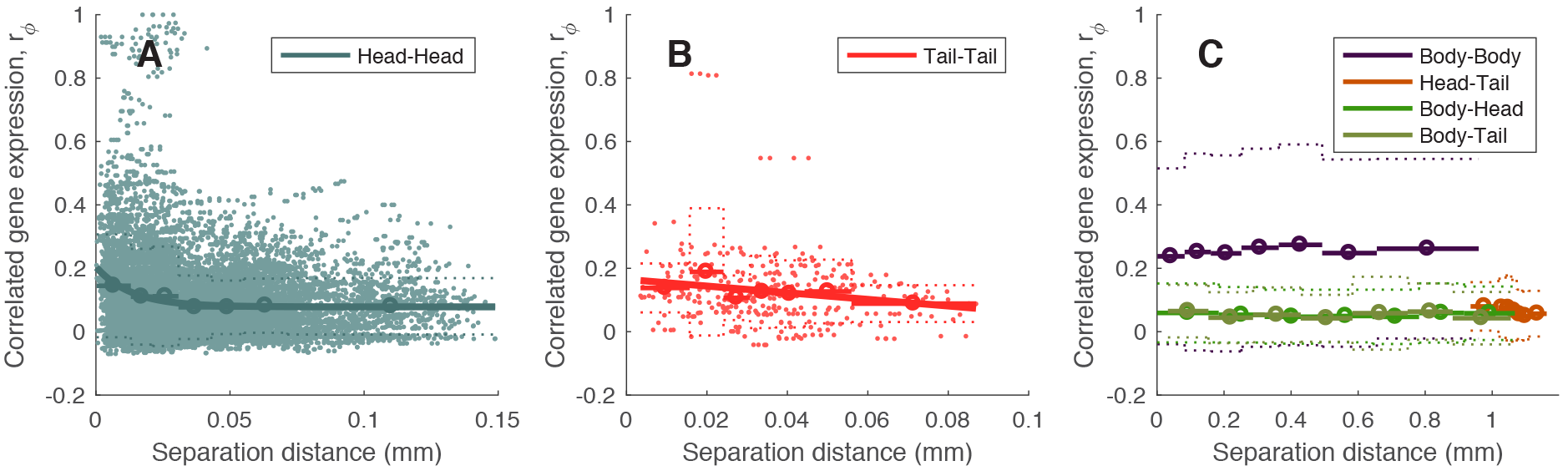
Dependence of correlated gene expression, *r*_*ϕ*_, on spatial separation between pairs of neurons. Correlated gene expression, *r*_*ϕ*_ (excluding bilateral homologous pairs of neurons), is shown as a function of the pairwise separation distance between pairs of neurons (shown as the mean (solid) ± standard deviation (dotted) in seven equiprobable distance bins, with extent shown as horizontal bars), for (A) all pairs of neurons in the head, (B) all pairs of neurons in the tail, and (C) all other pairs (labeled). Scatters for all neuron pairs are added in (A) and (B). An exponential relationship, *f*(*x*) = *A*exp(−*λx*) + *B*, is fitted in (A) and (B). The weak decreasing trend in *r*_*ϕ*_ with distance, is primarily driven by a small subset of nearby neurons with high *r*_*ϕ*_, and may therefore represent a more specific relationship between particular neurons, rather than a general, bulk spatial relationship observed in macroscopic mammalian brains [38, 72].

**Fig 5.**
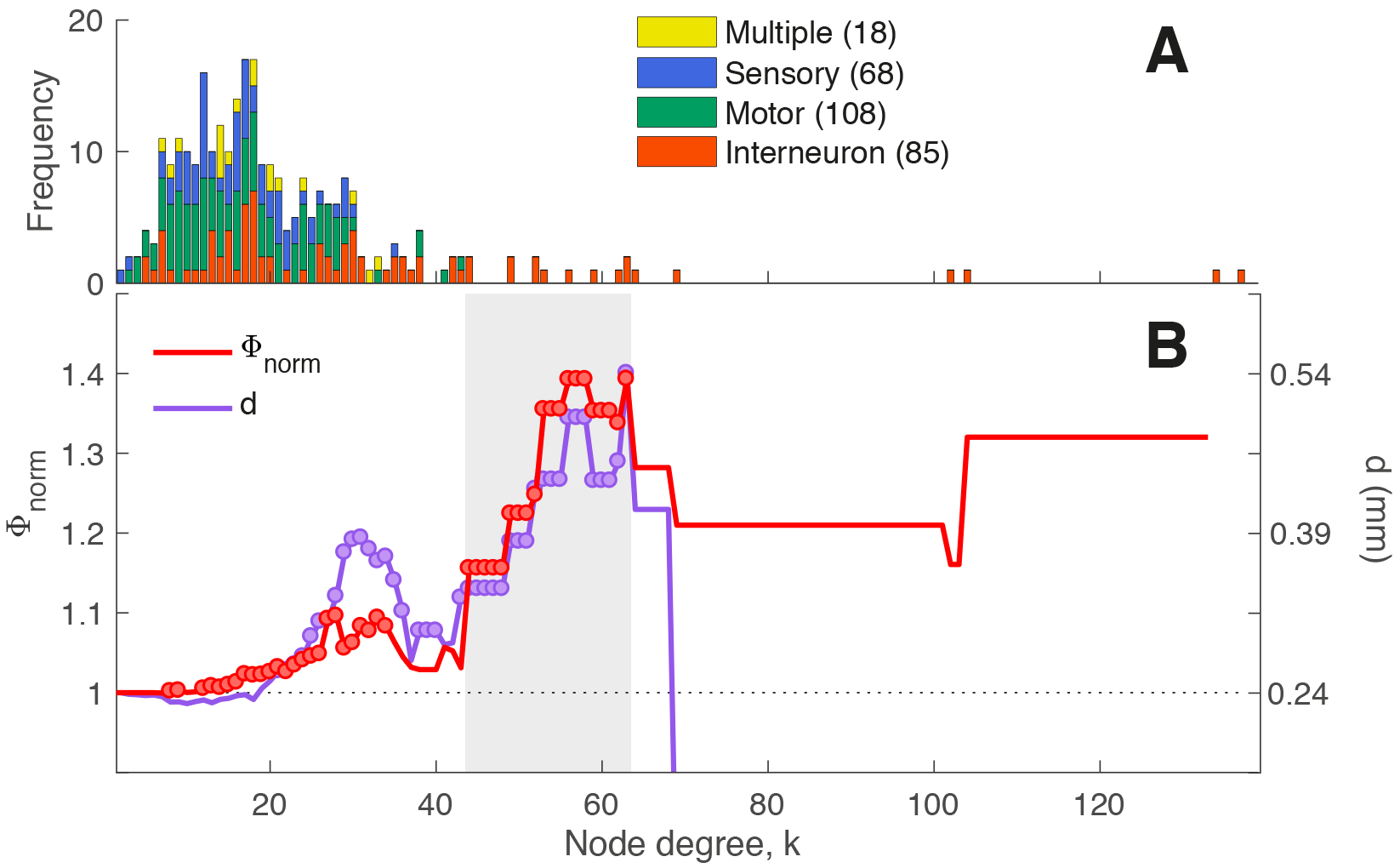
Rich-club organization of the *C. elegans* connectome. (A) Degree distribution of neurons, labeled to four categories: (i) interneuron (85 neurons, orange), (ii) motor (108 neurons, green), (iii) sensory (68 neurons, blue), or (iv) multiple assignments (18 neurons, yellow). The distribution features an extended tail of high-degree interneurons. (B) Normalized rich club coefficient, Φ_norm_ (red), as a function of the degree, *k*, at which hubs are defined (as neurons with degree > *k*). Also shown is the mean Euclidean separation distance, *d* (purple) between connected hub regions (across degree thresholds, *k*). Φ_norm_ > 1 indicates that hubs are more densely interconnected among each other than expected by chance, with red circles indicate values of Φ_norm_ that are significantly higher than an ensemble of 1 000 degree-matched null networks (*p* < 0.05). Purple circles indicate where the Euclidean distance between connected pairs of hubs is significantly greater than the Euclidean distance for all other pairs of connected regions (right-tailed Welch’s *t*-test, *p* < 0.05).

### Correlated gene expression and connectivity

We next investigate how the network connectivity properties of *C. elegans* relate to patterns of CGE, using the mean square contingency coefficient, *r*_*ϕ*_. To test whether CGE varies as a function of connectivity, we compared the distribution of *r*_*ϕ*_ between (i) all connected pairs of neurons, and (ii) all unconnected pairs of neurons. Connected pairs of neurons have more similar expression profiles than unconnected pairs (Wilcoxon rank-sum test, *p* = 1.8 × 10^−78^), mirroring previous results in the mesoscale mouse connectome [38]. Figure 6A (left) shows distributions of *r*_*ϕ*_ for: (i) all pairs of neurons that are connected via an electrical gap junctions (474 pairs, after excluding bilateral pairs), (ii) all pairs of neurons that are connected via reciprocal (291 pairs) and, (iii) unidirectional chemical synapses (1721 pairs) as well as (iv) all pairs of neurons that have neither connection (36 450 pairs). Note that 175 pairs of neurons are connected both by a chemical synapse and by a gap junction, and are thus included in both chemical and electrical categories. Amongst connected pairs of neurons, those connected via gap junctions exhibit more similar gene expression profiles than those connected via chemical synapses (Wilcoxon rank-sum test, *p* = 5.4 × 10^−22^). We found no difference in CGE between pairs of neurons connected reciprocally by chemical synapses (*N*_1_ ↔ *N*_2_ for two neurons *N*_1_ and *N*_2_) versus those connected unidirectionally (*N*_1_ → *N*_2_) (Wilcoxon rank-sum test, *p* = 0.99), in contrast to significant differences found in the mesoscale mouse connectome [38].

We next investigated whether CGE varies across different types of connections defined in terms of their hub connectivity. For a given hub threshold, *k*, we first labeled each neuron as either a ‘hub’ (nodes with degree > *k*) or a ‘nonhub’ (degree ≤ *k*), and then labeled each connection as either ‘rich’ (hub → hub), ‘feed-in’ (nonhub → hub), ‘feed-out’ (hub → nonhub), or ‘peripheral’ (nonhub → nonhub). The median CGE, *r̃_ϕ_*, of each of these four connection types is plotted in Fig. 6B, with circles indicating statistically significant increases of a given connection type relative to all other connections (one-sided Wilcoxon rank-sum test, *p* < 0.05). Correlated gene expression in rich connections increases with degree, *k*, particularly in the topological rich-club regime where hubs are densely interconnected (shaded gray in Fig. 6B). In this topological rich-club regime, both feed-in and feed-out connections exhibit increased CGE relative to peripheral connections, which show the lowest levels of CGE. Full distributions of *r*_*ϕ*_ for each edge type at a hub threshold of *k* > 44 are in Fig. 6A (right). This plot shows that, compared to all different types of pairs of neurons, connected pairs of hubs showed the highest CGE.

**Fig 6.**
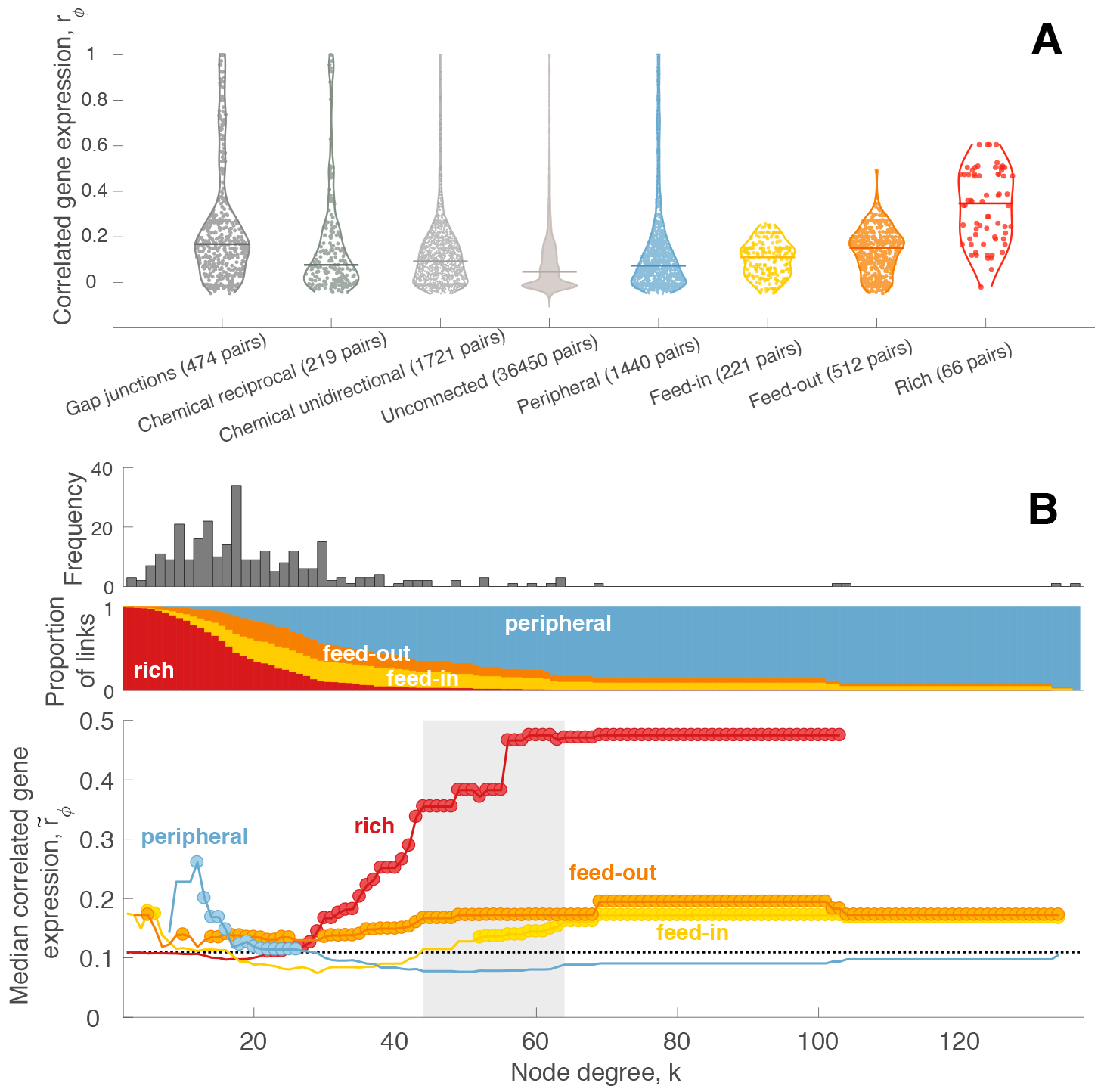
Correlated gene expression varies as a function of connectedness and connection type. (A) *Left*: distribution of CGE for (i) pairs of neurons connected by gap junctions, (ii) pairs of neurons connected by reciprocal chemical synapses, (iii) pairs of neurons connected by unidirectional chemical synapses, (iv) pairs of neurons that are unconnected, shown as a violin plot, with the median of each distribution represented by a horizontal line. CGE is increased in connected (electrical or chemical; reciprocally or unidirectionally) pairs of neurons relative to unconnected pairs (*p* = 1.8 × 10^−78^, Wilcoxon rank sum test). Among connected pairs of neurons neurons connected via gap junctions have more similar CGE than connected via chemical synapses (Wilcoxon rank-sum test, *p* = 5.4 × 10^−22^). *Right*: GCE for pairs of neurons labeled as peripheral, feed-in, feed-out, and rich, where hubs are neurons with degree *k* > 44. The median of each distribution shown as a horizontal line. CGE is significantly higher between hubs (rich links) compared to feeder (*p* = 5 × 10^−22^, Wilcoxon rank sum test) and peripheral (*p* = 3.9 × 10^−19^, Wilcoxon rank sum test) links. Feed-out links show significantly higher CGE than both feed-in (*p* = 1.9 × 10^−6^, Wilcoxon rank sum test) and peripheral links (*p* = 4.5 × 10^−12^, Wilcoxon rank sum test). (B) *Top*: Degree distribution, *k*, of the *C. elegans* connectome. *Middle*: proportion of connections that are: ‘rich’ (hub → hub, red), ‘feed-in’ (nonhub→hub, yellow), ‘feed-out’ (hub→nonhub, orange), or ‘peripheral’ (nonhub→nonhub, blue) as a function of the degree threshold, *k*, used to define hubs. Note that at high *k* most neurons are labeled as nonhubs and hence the vast majority of connections are labeled ‘peripheral’. *Bottom*: Median CGE, *r̃_ϕ_*, for each connection type as a function of *k*. The median CGE across all network links is shown as a dotted black line; the topological rich-club regime (determined from the network topology, cf. Fig. 5) is shaded gray. Circles indicate a statistically significant increase in CGE in a given link type relative to the rest of the network (one-sided Wilcoxon rank-sum test, *p* < 0.05).

In summary, our results reveal: (i) increased CGE in connected pairs of neurons; (ii) the highest CGE in rich connections; and (iii) lowest CGE in peripheral connections. These results, obtained using incomplete binary annotations of gene expression across 948 genes in a microscale neuronal connectome, are consistent with a prior analysis of the expression of over 17 000 genes across 213 regions of the mesoscale mouse connectome [38].

### Potential drivers of elevated correlated gene expression between hubs

The sixteen hub neurons in *C. elegans* (*k* > 44) are all interneurons; are all located in either the head or tail; are mostly contained within a single topological module of the network; are mostly cholinergic (13/16); and are all born prior to hatching. We therefore investigated whether the similarity of gene expression profiles between hubs is specific to their high levels of connectivity, or whether it could instead be driven by these other characteristics.

#### Interneurons

The sixteen hubs in *C. elegans* are all interneurons. To determine whether the increase in CGE in rich connections was specific to interneurons, we plotted the median CGE for hub-hub connections, *r̃_ϕ_*, as a function of the degree threshold, *k*, separately for connections involving interneurons, sensory neurons, and motor neurons, as shown in Figs. 7B. For the curve labeled ‘sensory’, for example, each point is the median *r̃_ϕ_* across connections involving sensory neurons (i.e., at least one neuron of a connected pair is a sensory neuron), for which both neurons have degree > *k*. The increase in median hub-hub CGE is strongest for connections involving interneurons. Motor neurons show a smaller increase with *k*, although the absence of motor and sensory neurons with high *k* makes it difficult to draw firm conclusions. However, we do find that CGE is higher for hub-hub pairs of interneurons compared to connections between all pairs of nonhub interneurons (Wilcoxon rank sum test, *p* = 5 × 10^−21^), indicating that the high CGE of rich pairs cannot simply be explained by the fact that all hub neurons are interneurons (Fig. 7C). We next investigated whether greater CGE of hub-hub pairs of neurons could be driven by specific anatomical properties of hub interneurons. Specifically, we selected a subset of nonhub interneurons that most closely resemble the anatomical properties of hub interneurons in terms of their position and projection pattern; that is, the cells are in similar locations in the head and their axons project to similar targets in the tail. These neurons were AVFL, AVFR, AVHL, AVHR, AVKR, AVJL, and AVJR. Pairs of hub interneurons show higher median CGE than pairs of anatomically-matched nonhub interneurons, with the difference being at the threshold of statistical significance (Wilcoxon rank sum test, *p* = 0.051), suggesting that the increase in CGE amongst hub interneurons cannot be explained by their anatomical similarity among interneurons.

#### Modular organization

Another important topological property of neural systems is modularity, whereby network nodes coalesce into tightly connected subsystems that are thought to serve a common function [84]. Prior work in humans has shown that functional networks in the brain have elevated transcriptional coupling [85]. The *C. elegans* connectome has a modular organization, with prior work decomposing it into: (i) modules of neurons with dense intra-module connectivity (and relatively sparse connectivity between modules) [22, 51, 54], or (ii) groups of neurons with more similar connectivity patterns within groups than between groups [52, 64]. We therefore examined the association between topological modularity of the *C. elegans* connectome and CGE. We used the Louvain community detection algorithm [65] to extract modules from the *C. elegans* connectome using consensus clustering (see *Methods*). Four modules were extracted, with eleven hubs in module one (which contains 111 neurons), four hubs in module two (96 neurons), one hub in module three (40 neurons), and no hubs in module four (32 neurons). We also compared the results of this modular partition of neurons to a previously reported nine-module partition derived from an Erdös-Rényi Mixture Model (ERMM) [52]. For the Louvain consensus modules, CGE, *r*_*ϕ*_, was significantly increased for connected neurons in the same module (1552 pairs) relative to connected pairs in different modules (687 pairs) (Wilcoxon rank sum test, *p* = 6.6 × 10^−4^), but there was no significant difference between intra-modular connected neurons and inter-modular connected neurons for the nine-module ERMM partition (Wilcoxon rank sum test, *p* = 0.46). The resolution and type of modular decomposition thus affects the relationship between CGE, connectivity, and modular network structure in *C. elegans*. We then tested whether connected hubs exhibit more similar CGE within and between modules (for both the consensus Louvain and ERMM modular decompositions). For pairs of connected neurons within the same module, *r*_*ϕ*_ is higher for pairs of hubs than pairs of nonhubs (Wilcoxon rank sum test, *p* = 6.9 × 10^−17^ for consensus Louvain modules, shown in Fig. 8A; 9.3 × 10^−7^ for ERMM partition). We found a similar result for connected neurons in different modules: pairs of connected hubs exhibit increased CGE than other types of connected pairs of neurons (Wilcoxon rank sum test, *p* = 1.6 × 10^−5^ for consensus Louvain modules, shown in Fig. 8B; *p* = 1.6 × 10^−16^ for ERMM). Thus, for both types of modular decompositions considered, intra-modular and inter-modular connections involving hub neurons exhibit more correlated gene expression patterns than other intra-modular and inter-modular connections.

**Fig 7.**
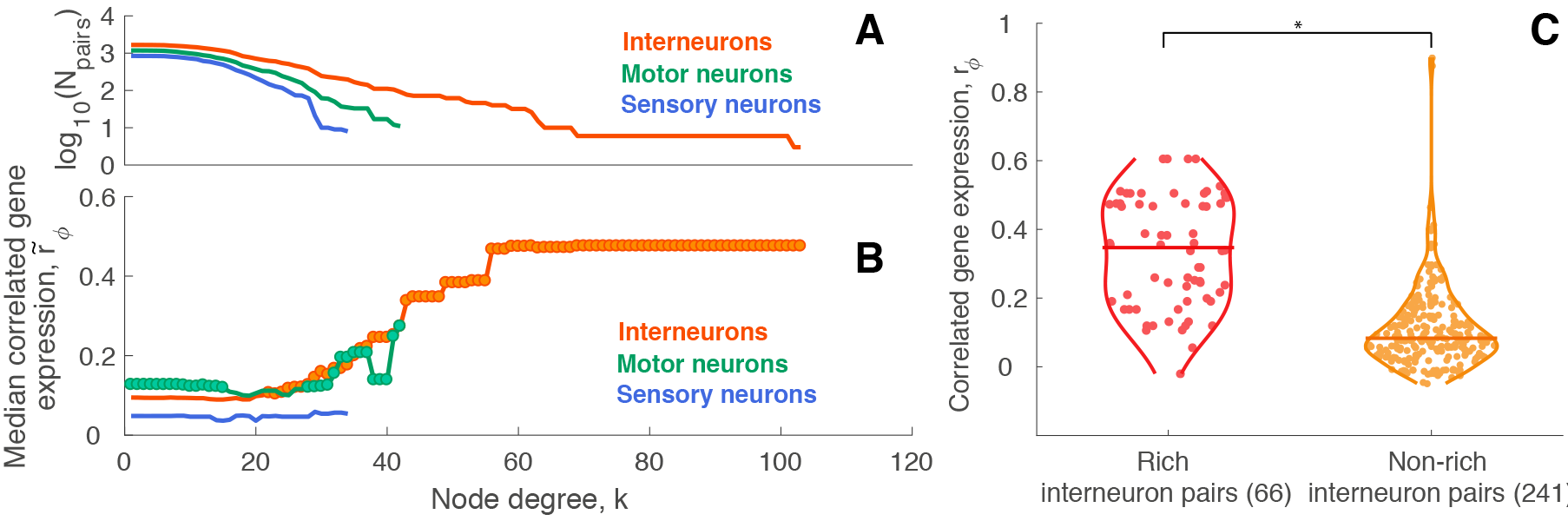
Correlated gene expression is highest for hub interneurons. (A) The number of connected neuron pairs involving interneurons (orange), sensory neurons (blue), and motor neurons (green) across degree threshold, *k*, represented as log10(number of links). (B) Median CGE as a function of degree for connections involving different types of neurons. Circles indicate a statistically significant increase in CGE in a given link type relative to the rest of the network (one-sided Wilcoxon rank-sum test, *p* < 0.05). (C) CGE distributions for connected pairs of hub interneurons (red) and connected pairs of non-hub interneurons (dark yellow) (Wilcoxon rank sum test, *p* = 5 × 10^−21^). * represents statistically significant difference.

#### Lineage distance

The lineage distance between a pair of neurons is defined as the sum of total divisions that have taken place since the most recent common ancestor cell [52, 59, 60]. In the mammalian brain, neuronal lineage has been associated with both functional properties [86, 87] as well as connectivity [88]. Moreover, tissue distance (resembling lineage distance on a cellular scale) correlates with gene expression divergence, meaning that tissues from the same branch on the fate map share more similar gene expression patterns in both human and mouse mesoderm as well as ectoderm tissues [89]. Given that the ectoderm eventually differentiates to form the nervous system, this finding suggests a possible relationship between lineage distance and CGE in a microscale neuronal system such as that of *C. elegans*. However, we find no significant correlation between lineage distance and CGE in *C. elegans* (Spearman’s *ρ* = 0.027, *p* = 0.2). As shown in Fig. 8C, there was only a weak tendency for the meadian lineage distance lineage distance to be increased in non-rich pairs (Wilcoxon rank sum test, *p* = 0.079). Thus, we can not attribute the transcriptional similarity of connected hub neurons to their neuronal lineage.

**Fig 8.**
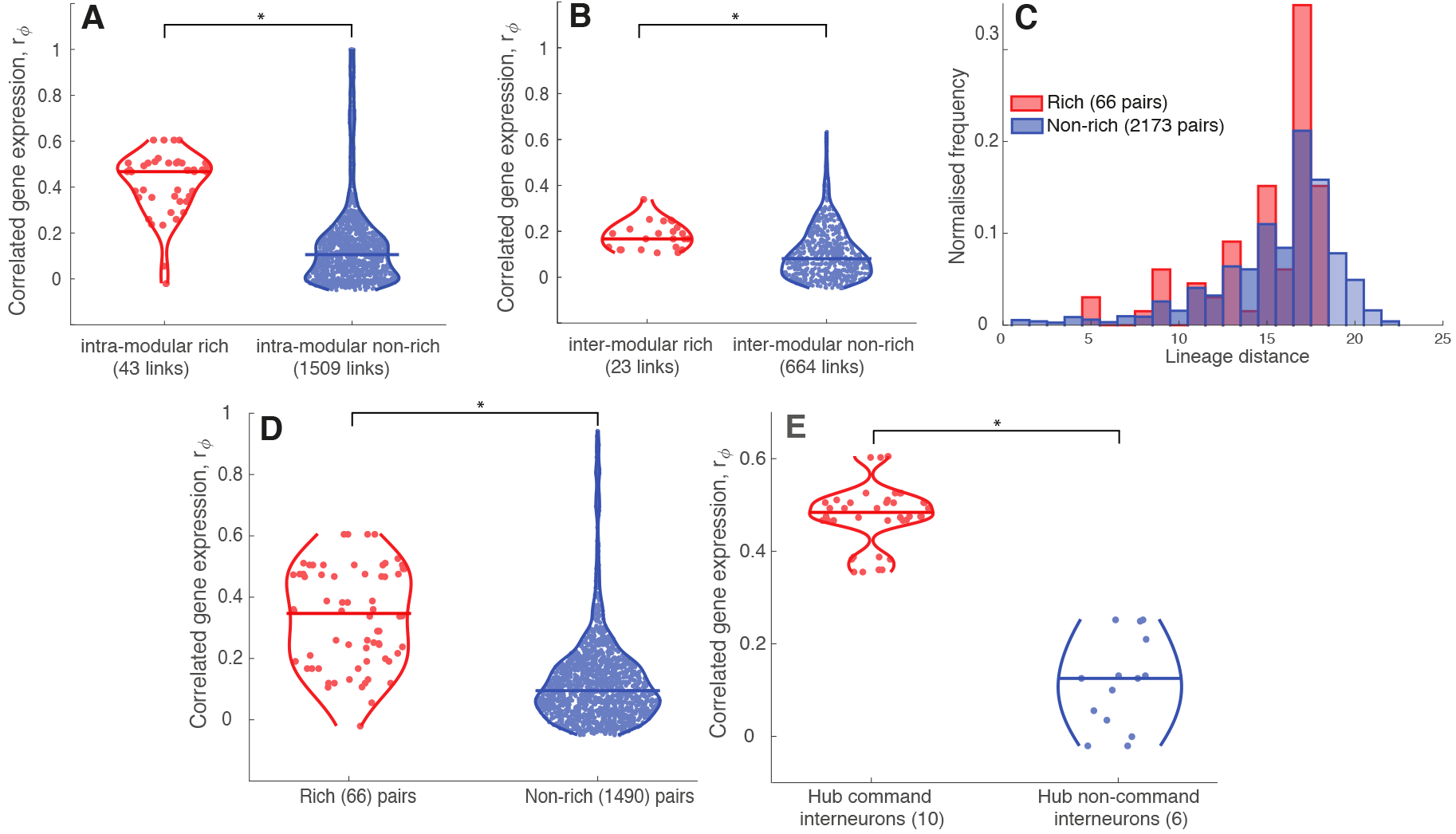
Increased CGE of hub neurons is not driven by modularity, neuronal birth time, or cell lineage distance. (A) Distributions of CGE, *r*_*ϕ*_, for intra-modular rich (red) non-rich (blue) connections, shown as violin plots with the median shown as a horizontal bar (Wilcoxon rank sum test, *p* = 6.9 × 10^−17^). (B) Distributions of CGE, *r*_*ϕ*_, for inter-modular rich (red) and non-rich (blue) connections, shown as violin plots with the median shown as a horizontal bar (Wilcoxon rank sum test, *p* = 1.6 × 10^−5^). (C) Distributions of lineage distance between rich links (red) and non-rich links (blue), plotted as histograms due to a discrete nature of this measure (Wilcoxon rank sum test, *p* = 0.079). (D) Distributions of CGE, *r*_*ϕ*_, between early born hubs (rich links, red) and nonhubs (non-rich links, blue) shown as violin plots with the median shown as a horizontal bar (Wilcoxon rank sum test, *p* = 3.9 × 10^−22^). (E) Distributions of CGE between hub command interneurons (red) and hub non-command interneurons (blue) shown as violin plots with the median shown as a horizontal bar (Wilcoxon rank sum test, *p* = 3.3 × 10^−8^). * represents statistically significant differences.

#### Birth time

The genesis of neurons in *C. elegans* is separated into two distinct time periods: before hatching (birth time < 550 min – ‘early-born’) and after hatching (birth time > 1200 min – ‘late-born’), with no neurons formed during intermediate times [56]. As a broad group, connected pairs of early-born neurons do not exhibit significantly different CGE compared to connected pairs of late-born neurons (Wilcoxon rank sum test, *p* = 0.64), but connected pairs of early-born neurons do exhibit significantly higher CGE relative to pairs of connected neurons for which one neuron is born prior to hatching and the other is born after hatching (Wilcoxon rank sum test, *p* = 4.2 × 10^−3^). Since all *C. elegans* hub neurons are born prior to hatching [28] and neurons born at similar times may share similar connectivity properties [28, 56], we investigated whether the increase in CGE between hub neurons of *C. elegans* may be driven by their similar birth times. Focusing on the 201 neurons born prior to hatching (16 hub and 185 nonhub neurons), connected pairs of hub neurons exhibit significantly increased CGE, *r*_*ϕ*_, relative to other pairs of connected early-born neurons (Wilcoxon rank sum test, *p* < 10^−22^), as shown in Fig. 8D.

#### Neurochemistry

Hub neurons (*k* > 44) consist of thirteen cholinergic neurons, two glutamatergic neurons, and one neuron of unknown neurotransmitter type [58]. We find that neuron pairs show different CGE relationships as a function of their neurotransmitter type, e.g., pairs of GABAergic neurons have high median CGE, *r̃_ϕ_* = 0.59, while pairs of glutamatergic neurons exhibit a relatively low median CGE, *r̃_ϕ_* = 0.08. To determine whether the similarity in CGE between pairs of hub neurons could be explained by their neurotransmitter types, we constructed 10^8^ random sets of sixteen neurons of the same neurotransmitter types as hubs (e.g., thirteen random cholinergic neurons, two random glutamatergic neurons, and one random neuron of an unknown neurotransmitter type) and compared the distribution of median CGE of each group to the median CGE of hub neurons as a permutation test. Neurochemical identity cannot explain the elevated CGE of hub neurons, with hubs displaying a significant increase in median CGE relative to random sets of neurons with the same neurotransmitter types as hubs (permutation test, *p* = 2.57 × 10^−6^).

#### Anatomical location

Of the sixteen hub neurons, thirteen are in the head and three are in the tail; none are located in the body. CGE varies as a function of anatomical location (i.e., ‘head’, ‘body’, and ‘tail’), with pairs of neurons in the same anatomical class exhibiting the highest median CGE (e.g., pairs of neurons within the body have a median *r̃_ϕ_* = 0.14, pairs of neurons within the tail have *r̃_ϕ_* = 0.12, and pairs of neurons within the head have *r̃_ϕ_* = 0.07), followed by lower median CGE in mixed classes (e.g., head-body pairs of neurons have *r̃_ϕ_* = 0.01). Given this variation, we tested whether the increased CGE between hub neurons could be explained by their anatomical distribution using the permutation testing procedure described above for neurochemistry. That is, we compared the distribution of CGE between hubs to a null distribution formed from 10^8^ random permutations of thirteen head neurons and three tail neurons. The median CGE, *r̃_ϕ_*, between hub neurons is significantly increased relative to random sets of thirteen head neurons and three tail neurons (permutation test, *p* = 2.62 × 10^−6^). Furthermore, CGE is significantly increased amongst hub neurons relative to other pairs of head neurons (Wilcoxon rank sum test, *p* = 4.1 × 10^−11^ for 46 hub-hub pairs and 1 186 others), and also amongst head/tail pairs of neurons (Wilcoxon rank sum test, *p* = 1.6 × 10^−14^ for 23 hub-hub pairs, 174 others), but not for the three hub neurons in the tail (Wilcoxon rank sum test, *p* = 0.15 for three hub-hub pairs, 53 others). Thus, anatomical location cannot explain the high CGE of hub neurons.

#### Functional class

*C. elegans* neurons can be divided into distinct groups that each perform a specialized behavioral function [90]. One of the best-characterised functional classes that is particularly relevant for our analysis is the set of ‘command interneurons’ which are a functional group of ten neurons that govern forward (AVBL, AVBR, PVCL, PVCR) and backward (AVAL, AVAR, AVDL, AVDR, AVEL, AVER) locomotion [91]. All of these neurons are hubs. Given the overlap between hub neurons and command interneurons, we investigated whether command interneurons exhibit more similar expression than other hub interneurons, and may therefore drive the increase in CGE amongst hubs as a whole. We compared CGE, *r*_*ϕ*_, between all pairs of command interneurons (ten neurons), and between all pairs of hubs that are not command interneurons (six neurons), shown in Fig. 8E. Correlated gene expression between command interneurons is significantly greater than between other hub neurons (Wilcoxon rank sum test, *p* = 3.3 × 10^−8^), indicating that command interneurons play a major role in driving the increased CGE amongst hub neurons. Moreover, there is no difference in CGE between pairs of hubs that are not command interneurons (DVA, RIBL, RIBR, AIBR, RIGL, AVKL) and a set of seven anatomically matched nonhub head interneurons (AVFL, AVFR, AVHL, AVHR, AVJL, AVJR, AVKR) (Wilcoxon rank sum test, *p* = 0.13), indicating that the status of many hub neurons as command interneurons makes a significant contribution to the elevated CGE of hubs.

### Genes driving correlated gene expression patterns

Having characterized a robust relationship between CGE and (i) connectivity, and (ii) hub connectivity, we next investigated which specific genes contribute most to this relationship. Despite challenges with the incomplete binary expression measurements in a small proportion of the genome, we developed a method to score genes according to their contribution to a given CGE pattern (see *Methods*). We characterized individual high-scoring genes, with *p*_corr_ < 10^−4^ (approximately 20% of genes with the highest scores in each analysis), and attempted to summarize functional groups of genes as biological process categories of the gene ontology (GO) that were enriched in high scoring genes using overrepresentation analysis (ORA) [68, 70].

We first investigated which genes drive increased CGE in connected pairs of neurons relative to unconnected pairs. Previous studies in mouse have indicated that genes driving an increase in CGE between connected pairs of brain regions are enriched in GO categories related to neuronal connectivity and communication [38, 92, 93, 94]. First, we manually investigated individual high-scoring genes (i.e., those with *p*_corr_ < 10^−4^, see Supplementary Data File, **GeneList.xlsx**). Given that glutamate is a prevalent neurotransmitter in *C. elegans* (26% of neurons with known neurotransmitter type are glutamatergic [58]), it is appropriate that many high scoring genes are related to glutamate receptors (including *glr-1*, *glr-2*, *glr-4*, *glr-5*, *nmr-1*, and *nmr-2*). Consistent with the importance of innexins in forming electrical synapses [95], our list contained the following innexin genes: *unc-9*, *unc-7*, *inx-7*, *inx-19*, *inx-13*. Genes encoding cell adhesion molecules related to axon outgrowth and guidance, cell migration and locomotion (*sax-3*, *cam-1*, *unc-6*, *rig-1*, *unc-5*), learning (*casy-1*) [57, 96, 97, 98, 99], as well as genes involved in determining cell polarity (*vang-1*, *prkl-1*) [100, 101] were also amongst the top scoring genes for connectivity. These genes have been implicated in neuronal connectivity in both flies and humans [102, 103, 104], with our results predicting that they may play a similar role in *C. elegans*. In addition, transcription factors regulating neuronal development, fate specification (*lin-11*, *unc-3*, *unc-42*, *ceh-14*, *ast-1*, *cfi-1*) [105, 106, 107, 108, 109, 110], and locomotion (*unc-3*) [106]) were also implicated in driving increased CGE amongst connected pairs of neurons. Both adhesion molecules and transcription factors are candidates for facilitating signal transduction and communication.

In order to summarize the above mentioned results and determine if any particular functional groups of genes are dominating in driving this effect, we performed an enrichment analysis. Top scoring biological process GO categories from ORA analysis (of 85 genes relative to the 414 genes with sufficient data for this analysis) are listed in Table S2. Although no GO categories are significant at a false discovery rate of 0.05, the top categories are consistent with a connectivity profile, including ‘glutamate receptor signaling’, ‘cell surface receptor signaling’, and ‘ion transport’, with other categories involved in regulation of growth rate and several related to catablic processes. Thus, despite incomplete gene expression data that do not provide sufficient coverage to detect statistically significant effects, these results indicate that our data-driven gene scoring method is able to yield sensible, biologically relevant insights into the genetic basis of neuronal connectivity in *C. elegans* connectome.

Having characterized genes that contribute to the increase in CGE between connected pairs of neurons, we next investigated whether particular functional groups of genes drive differences in CGE between connections involving hub neurons (i.e., in rich, feed-in, and feed-out connections) relative to connections between pairs of nonhub neurons (i.e., peripheral connections). In order to investigate which specific genes contribute most to the increase in CGE for connections involving hubs, we first investigated
the highest-scoring genes, with *p*_corr_ < 10^−4^ (corresponding to approximately the top 20% of genes in the analysis). In addition to glutamate receptor genes (*glr-5*, *nmr-1*, *nmr-2*, *glr-1*, *glr-2*, *grld-1*) and acetylcholine related genes (*ace-2*, *cho-1*, *unc-17*, *deg-3*), we again find a high number of genes regulating cell adhesion (*cam-1*, *rig-1*, *rig-6*, *unc-6*, *grld-1*, *dbl-1*, *ncam-1*) and relevant transcription factors (*unc-3*, *unc-42*, *ast-1*). The implication of glutamate and acetylcholine may be attributable to the importance of glutamate in the regulation of locomotion in command interneurons [111, 112], with acetylcholine being the dominant neurotransmitter in hubs (13 out of 16 hubs are cholinergic). We also find a high overlap between adhesion molecule and transcription factor genes found in the previous analysis and the implication of human orthologs (*rig-1*, *ncam-1*, *grld-1*, corresponding to human genes ROBO4, NCAM2, and RBM15 respectively [57]) for genes regulating cell migration, differentiation and neuron cell adhesion.

While previous work implicated genes regulating oxidative metabolism for connections involving hubs in mouse [38], and for hub regions in human [44], the gene expression dataset used here was not sufficiently comprehensive to investigate these processes. For example, only one of the 948 genes annotated to the GO categories related to hub connectivity in mouse is present in our gene expression dataset (*unc-32* is annotated to the GO category: ‘ATP hydrolysis coupled proton transport’). Thus, although a direct test of the metabolic hypothesis for neural hubs is not possible from current data, we investigated whether other biological process GO categories were overrepresented in pairs of connected hubs using ORA (of 30 genes relative to the 168 genes with sufficient data for this analysis), with results listed in Table S3. Even though no categories are statistically significant at a false discovery rate of 0.05, the list of top categories includes both ‘glutamate receptor signaling pathway’ as well as more general ‘cell surface receptor signaling pathway’ in addition to several ion transport related gene groups (‘ion transport’, ‘ion transmembrane transport’, ‘transmembrane transport’). Other top-ranked GO categories include regulation of locomotion, and various metabolism and biosynthesis related processes. Our gene scoring method again yields interpretable insights into the types of genes that contribute to differences in CGE between different classes of neuronal connections in *C. elegans*. While current data are limited, more comprehensive expression annotations in the future would allow more systematic and statistically powered inferences across GO categories.

## Discussion

Highly connected hubs of neural systems play an important role in brain function, with their dense rich-club interconnectivity integrating disparate neural networks [24, 27, 29, 30]. Here, our analysis linking hub connectivity of the microscale connectome of *C. elegans* to patterns of neuron-specific gene expression has identified a transcriptional signature that appears to be highly conserved, given recent findings reported in a mesoscale investigation of the mouse [38] and a macroscale study of humans [44]. Specifically, we show that: (i) CGE is higher for connected pairs of neurons compared to unconnnected pairs; (ii) the neuron connection probability decays as a function of spatial separation, and; (iii) connected pairs of hub neurons, which are generally separated by longer anatomical distances, show the highest levels of CGE. This association between CGE and hub connectivity followed a gradient, such that CGE was lowest for connected nonhubs, intermediate for hub-nonhub pairs, and highest for connected hubs, consistent with results reported in the mouse brain [38]. Amongst the genes considered here, many of those with the greatest contribution to connectivity are biologically plausible genes related to receptors, neurotransmitters, and cell adhesion, and those with the greatest contribution to hub connectivity are related to glutamate receptors, acetylcholine signaling, and other neuronal communication related genes. The methods we develop here for quantifying CGE, and for scoring the contribution of individuals genes to overall CGE, yield biologically interpretable results from incomplete binary gene expression data. With improvements in gene annotation quality and specificity, and increases in genome coverage, similar methods could be used in future work to characterize the biological basis of a range of neuronal connectivity patterns.

It is reasonable to expect that the principles of neural organization may differ from the scale of individual neurons to the scale of macroscopic brain regions (in which each brain region contains millions of neurons). However, many of our results in *C. elegans* suggest a striking conservation of many fundamental spatial trends in neural connectivity and CGE across scales and species. For example, connection probability decreases with spatial separation between brain areas in rodents and primates [75, 76] (including in macaque [77], human [78], mouse [38], and rat [79]), for individual neurons in mouse primary auditory cortex [80], and between neurons in *C. elegans* (cf. Fig. S1 of [49]). Unlike mammalian brains, where all neurons are confined to a spatially contiguous organ, neurons are distributed across nearly the entire length of *C. elegans*, including a dense cluster of neurons in the head and in the tail. Despite these distinct morphologies, we report a qualitatively similar spatial dependence of connection probability with separation distance for many classes of connections in *C. elegans*, including those within the head, body, and tail, indicating that this distance-dependence may be a generic property of evolved neuronal systems that must balance the energetic cost of long-range connections with their functional benefit [17, 23, 33, 51]. Less frequently characterized is the spatial dependence of CGE, with available evidence indicating that more proximal brain areas exhibit more similar gene expression patterns than more distant brain areas in the mouse brain [38] and human cortex [72, 73, 74]. Some of the spatial trends in CGE found in the 948 genes analyzed here mirror these trends of bulk regions of macroscopic mammalian brains. It is therefore possible that these spatial dependences of connectivity and CGE may not be simply due to bulk spatial trends in macroscopic brains containing millions of neurons, but may reflect conserved organizational principles that hold across species and spatial scales. Our results highlight the importance of treating nervous systems as spatially embedded objects, as many seemingly non-trivial properties of brain organization may be well approximated by simple, isotropic spatial rules [22, 46, 75, 78, 113] (see also [17, 23]).

Our analysis indicates that CGE patterns in *C. elegans* show many surprising similarities to previous work in the mesoscale mouse connectome [38], despite: (i) involving different gene expression annotation data (comprehensive *in situ* hybridization expression data across ~ 20 000 genes in mouse versus literature-curated annotations across ~ 1000 genes in *C. elegans*), (ii) being a different type of neural system (from the spatially continuous macroscopic brain of mouse, to the spatially separated nervous system of *C. elegans*); (iii) orders of magnitude differences in spatial scale. The findings were also robust to a range of data processing choices, including different representations of the connectome (e.g., directed/undirected, or excluding electrical synapses), and across alternative metrics for quantifying transcriptional similarity.

What could drive this highly conserved association between CGE and hub connectivity? Here, we took advantage of the rich and diverse information available for each neuron of the *C. elegans* connectome to begin to address this question. We show that CGE between hub neurons cannot be explained by the neuronal subtype of hubs (i.e., the fact that all hubs are interneurons rather than sensory or motor neurons), since CGE between hub neurons is higher than between other pairs of interneurons. The effect also cannot be explained by the modular organization of the network, since CGE between hubs in the same module is higher than between other pairs of neurons in the same topological module, with a similar increase in CGE for pairs of hubs in different modules. We also show that the effect is not driven by similarities in the birth time nor lineage distance of hub neurons, which exhibit higher CGE than other early-born neurons (prior to hatching) and are not closer in their lineage. Moreover, the abundance of cholinergic signaling of hub neurons cannot explain the effect. Rather, the CGE between pairs of hub neurons in *C. elegans* may be related to the specific functional role of these cells. Namely, 60% of them are command interneurons, which play a vital role in coordinating forward and backward locomotion in *C. elegans* [50]. The overlap between command interneurons and hub neurons has interesting parallels with the human cortex, where polymodal association areas tend to be the most highly connected network elements [1]. Association areas sit at atop the cortical hierarchy and support complex behaviors by integrating information from diverse neural systems [114]. Locomotion is arguably one of the most complex behaviors expressed by *C. elegans*. Thus, the association between hub status and command interneurons may reflect the specialization of these neurons for supporting higher-order functions in the behavioral repertoire of *C. elegans*.

It is as yet unclear whether CGE between network hubs, regardless of species and scale, is simply a byproduct of tightly coupled hub activity, or some shared morphological or development characteristic between hubs that we have not captured in the present analysis. More comprehensive transcriptomic data (e.g., obtained through systematic single-neuron RNA sequencing), measured through development and coupled with measures of neuronal activity, would allow us to address these questions. Additionally, we cannot rule out the possibility that gene annotations have been influenced by the nature of the curated data that we have used here. Given their functional similarity, command interneurons might have been tested as a group in a set of experiments for the expression of particular genes and consequently assigned similar expression signatures. More precise and systematic measurement of neuron-specific gene expression patterns would be required to address this question.

In this work we developed methods to relate correlations in binary gene expression data to pairwise connectivity and subsequently score and evaluate the contribution of individual genes to these patterns. Compared to continuous *in situ* hybridization measurements of the expression of > 17 000 genes in the mouse brain [115], or microarray measurements of > 20 000 genes in the human brain [116, 117], which permit more detailed analysis [38, 44, 92, 93, 94, 118], working with *C. elegans* gene expression data is challenging due to its low coverage (< 5% coverage of the worm genome), binary indications of expression, and incompleteness (an inability to distinguish missing data from lack of expression). Moreover, the data have different qualifiers related to the certainty of gene expression annotations (see Supporting Information), requiring choices to be made to appropriately balance sensitivity and specificity. Although gene enrichment analyses did not have enough power to detect significant effects here, top GO categories point us towards biologically relevant categories related to neuronal connectivity, neurotransmitters, and metabolism. We note, however, that the incomplete coverage of the genome in our annotated dataset may mask many true GO associations. Our single gene analysis identified specific genes contributing to increases in CGE for connected pairs of neurons and for connections involving hub neurons. In line with our expectations, genes regulating both chemical and electrical signaling, namely glutamate receptor and innexin genes, were implicated in general connectivity. In addition, we also find multiple cell adhesion molecule genes and transcription factors that regulate neuronal development and fate specification – both groups are important for forming neuronal connections. High overlap between adhesion molecule genes and transcription factors implicated in regulating both general and hub connectivity highlights that related mechanisms might be used in both cases. While we were not able to test GO categories related to neurotransmitter signaling comprehensively, due to insufficient coverage of gene expression annotations, single gene analysis revealed the importance of acetylcholine genes, which may be related to the fact that acetylcholine is the dominant neurotransmitter in hub neurons.

## Acknowledgments

We thank WormBase, and particularly Daniela Raciti, for providing valuable information regarding the *C. elegans* gene expression data, and thank Taqi Ali for preliminary data processing. AF was supported by the Australian Research Council (ID: FT130100589) and the National Health and Medical Research Council (ID: 3251213). BDF was supported by an NHMRC Early Career Fellowship (1089718). RP was supported by Monash University Biomedicine Discovery Fellowship, NHMRC Project Grant (GNT1105374) and veski innovation fellowship: VIF 23 to RP.

## Supporting Information

### Expression annotations from WormBase

Neuronal gene expression is measured as a binary indicator on WormBase [57], based on curated data collated from many individual experiments. Expression annotations are made either ‘directly’ to individual neurons (when an experiment indicates expression in an individual neuron), or ‘indirectly’ to broader classes of neurons like ‘interneuron’ or ‘head’ (meaning that some members of that class exhibit expression of that gene). In order to maintain specificity of annotations, we only analyzed ‘direct’ annotations here.

Annotations of gene *G* to neuron *Y* are also made with varying levels of certainty:

- ‘certain’: *G* was observed to be expressed in *Y*,
- ‘enriched’: *G* has been found to be enriched in a certain dataset through microarray, RNAseq or proteomic analysis,
- ‘partial’: *G* was observed to be expressed in some cells of a group of neurons that include *Y*,
- ‘blank’: data prior to 2005, or
- ‘uncertain’: *G* was sometimes observed to be expressed in *Y*, or *G* was observed to be expressed in a cell that could be *Y*.

To avoid including false positives, our analysis excluded annotations labeled as ‘uncertain’.

### Correlated gene expression matching index

Existing measures of binary correlation (described below) are symmetric between ‘0’ and ‘1’ and thus do not directly distinguish between the biologically relevant case of genes being expressed together from the case in which neither gene is expressed. We thus developed a measure of the probability, *P* (*m*), that two binary strings will contain *m ‘*positive matches’ (i.e., *m* genes are expressed in both neurons), which can be computed as:

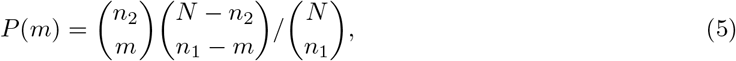

for two binary expression vectors, *x_i_*, *y_i_*, of length *N*, containing *n*_1_ and *n*_2_ 1s, respectively (*n*_2_ ≤ *n*_1_), with *m* matches (*n*_11_). Our index, *p*_match_, is approximately equal to the probability of obtaining as many or fewer matches than observed (weighting the probability of the observed number of matches at 0.5 for symmetry), computed as:

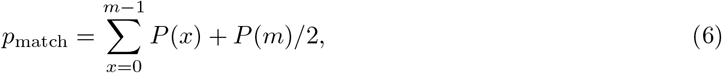

for *m* positive matches.

We verified that this index yields qualitatively similar results to the mean square contingency coefficient, *r*_*ϕ*_, used throughout this work (Fig. S2), validating our related positive match method for scoring individual genes [as an approximately single-gene contribution to the probability score computed in Eq. (6)]. Given the qualitative similarity of this measure to *r*_*ϕ*_, we chose to focus on *r*_*ϕ*_ throughout this work due to its ease of interpretability as a correlation coefficient ranging from -1 to 1.

### Sensitivity of correlated gene expression measures

Measures of correlation between binary vectors can be biased by the relative proportion of ones between two vectors. For data analyzed here, data annotated to individual neurons comes from between 3 to 138 expressed genes (corresponding to 0.3% to 14.6% of all 948 genes considered). To ensure that our measure of correlated gene expression (CGE) is not biased by such differences in the relative proportions of expression annotations, we conducted a sensitivity analysis in which we compared the *r*_*ϕ*_ metric, Eq. (3), with alternative methods for quantifying correlations between binary vectors: the Jaccard index, *n*_11_/(*n*_10_ + *n*_01_ + *n*_11_), Yule’s *Q* coefficient, (*n*_00_*n*_11_ − *n*_01_*n*_10_)/(*n*_00_*n*_11_ + *n*_01_*n*_10_), and the *χ*^2^ index, *N* (*n*_00_*n*_11_ − *n*_01_*n*_10_)/(*n*_1•_*n*_0•_*n*_•_*n*_•1_) [119], where *n_xy_* counts the number of observations of each of the four binary pairwise possibilities: *n*_00_, *n*_01_, *n*_10_, and *n*_11_ (as outlined in the main text), and the • symbol sums across a given variable (e.g., *n*_•0_ = *n*_00_ + *n*_10_).

To evaluate bias in each CGE measure to the proportion of annotations in each expression vector, we generated random binary vectors of length 948 containing different proportions of 1s seen in our data, ranging from the minimum, 1, to the maximum, 150. For all pairwise combinations of proportions, we computed the CGE measure, taking an average across 1 000 permutations, and then recorded the resulting mean correlation value, as plotted in Fig. S1. Because all vectors are independent random binary strings, any systematic dependence of mean CGE with annotation proportion indicates bias. The mean square contingency coefficient, *r*_*ϕ*_ (Fig. S1A) and our own novel CGE matching index, *p*_match_ (Fig. S1D) show no systematic dependence on the proportion of ones in each vector (varying randomly within ≈ 10^−3^ and ≈ 0.5 respectively). However, Yule’s *Q* shows a negative bias for small annotation proportions (Fig. S1B) and the Jaccard index shows a strong positive bias across the full range (Fig. S1C). Based on these numerical experiments, we selected mean square contingency coefficient, *r*_*ϕ*_, here, to ensure that changes in CGE were due to matching expression patterns and not simply driven by differences in the number of gene annotations between neurons.

**Table S1.**
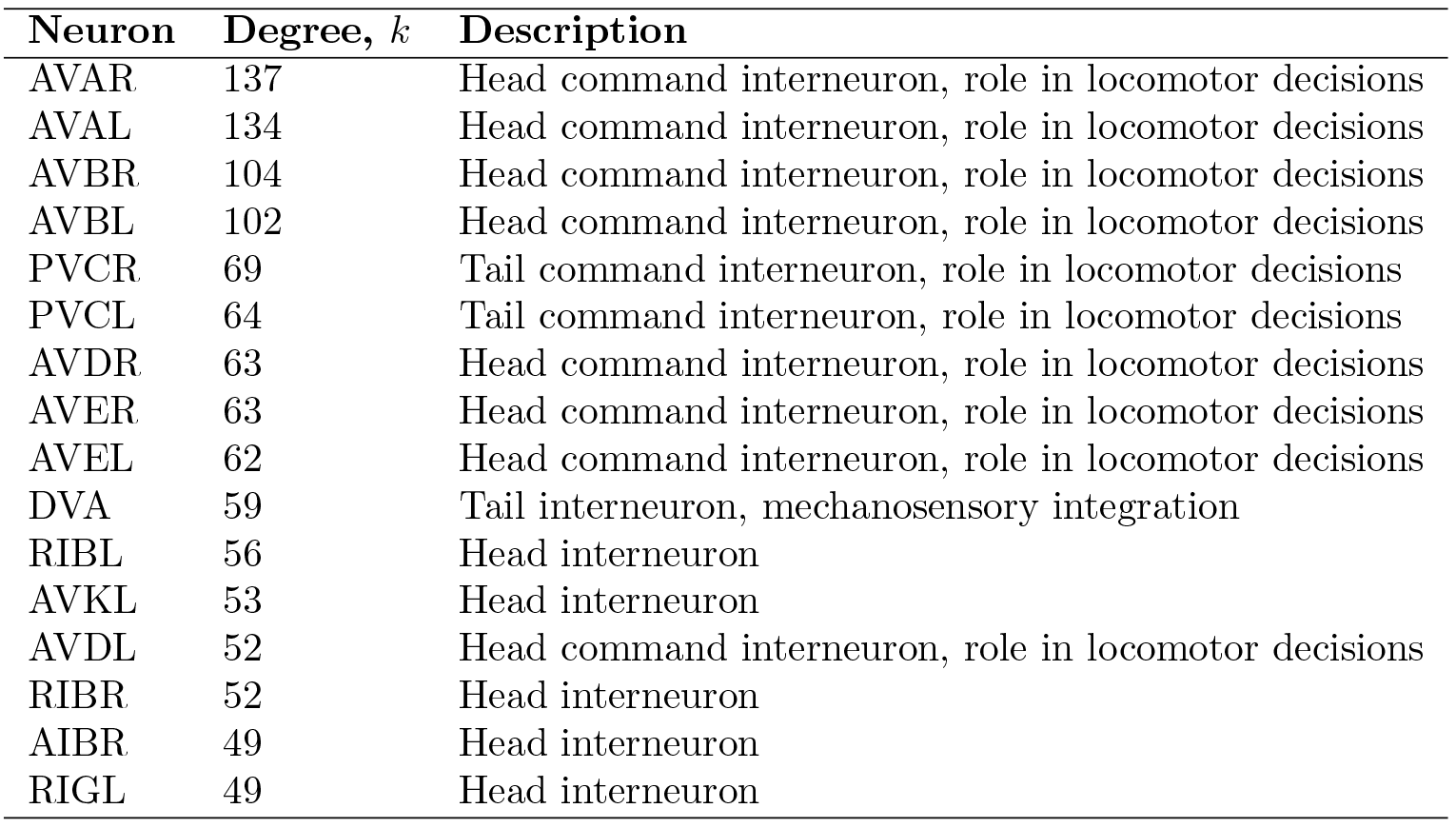
Hub neurons of the *C. elegans* connectome. Hubs are defined as neurons with degree *k* > 44. For each hub, we list (i) the neuron name, (ii) its degree, *k*, (iii) location (‘head’, ‘body’, or ‘tail’), and function based on information presented in the Wormatlas website, **www.wormatlas.org/**. Neurons are sorted (descending) by degree.

**Table S2.** Top 15 biological process GO categories enriched in genes with the highest mean increase in CGE for connected neurons compared to unconnected neurons. Categories are sorted by *p*-value (ascending).

| Category | Description | # genes | $p$ (uncorr) | $p$ (corr) |
| --- | --- | --- | --- | --- |
| GO:0035235 | ionotropic glutamate receptor signaling pathway | 7 | 0.0005 | 0.3245 |
| GO:0007215 | glutamate receptor signaling pathway | 9 | 0.0041 | 0.9367 |
| GO:0007166 | cell surface receptor signaling pathway | 37 | 0.0048 | 0.9367 |
| GO:0006811 | ion transport | 71 | 0.0058 | 0.9367 |
| GO:0034220 | ion transmembrane transport | 57 | 0.0079 | 1 |
| GO:0055085 | transmembrane transport | 67 | 0.0129 | 1 |
| GO:1901575 | organic substance catabolic process | 33 | 0.031 | 1 |
| GO:0040009 | regulation of growth rate | 7 | 0.0417 | 1 |
| GO:0040010 | positive regulation of growth rate | 7 | 0.0417 | 1 |
| GO:0030163 | protein catabolic process | 27 | 0.0428 | 1 |
| GO:0009056 | catabolic process | 35 | 0.0493 | 1 |
| GO:0009057 | macromolecule catabolic process | 28 | 0.0548 | 1 |
| GO:0071495 | cellular response to endogenous stimulus | 14 | 0.0575 | 1 |
| GO:0045927 | positive regulation of growth | 29 | 0.0689 | 1 |
| GO:0050830 | defense response to Gram-positive bacterium | 5 | 0.0695 | 1 |

**Table S3.**
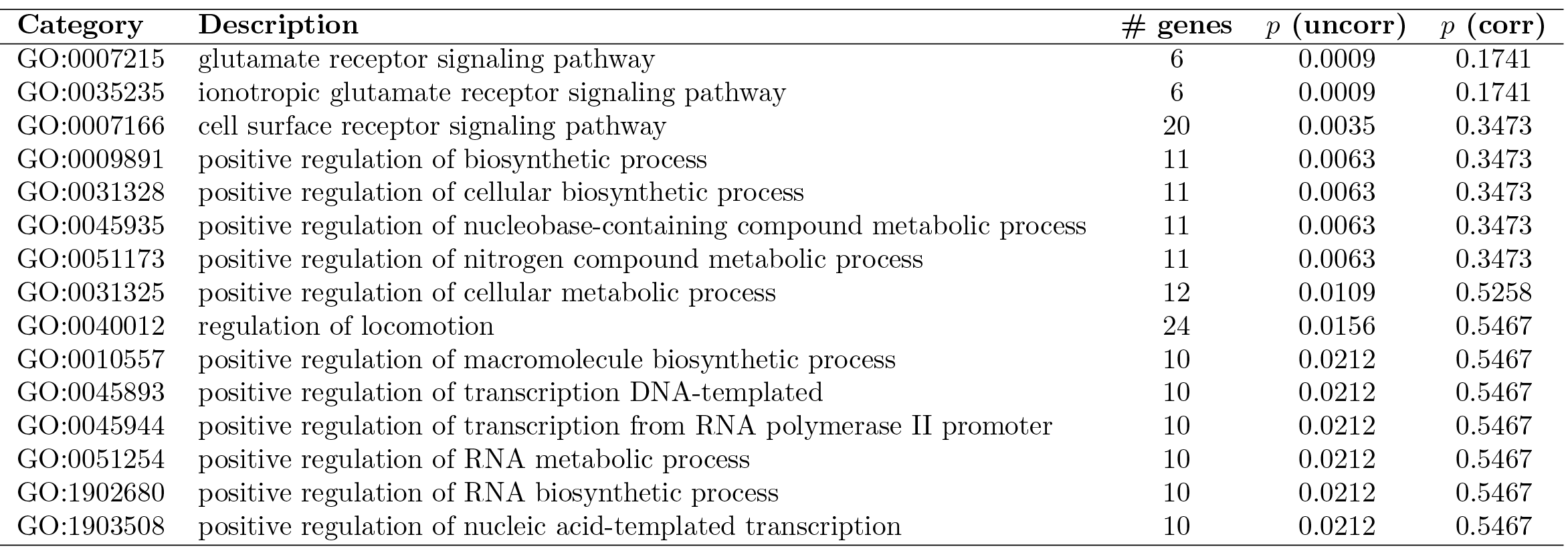
Top 15 biological process GO categories enriched in genes with the highest increase in CGE for connections involving hub neurons (i.e., rich, feed-in and feed-out connections) compared to connections between nonhub neurons (i.e., in peripheral connections). Categories are sorted by *p*-value (ascending).

**Fig S1.**
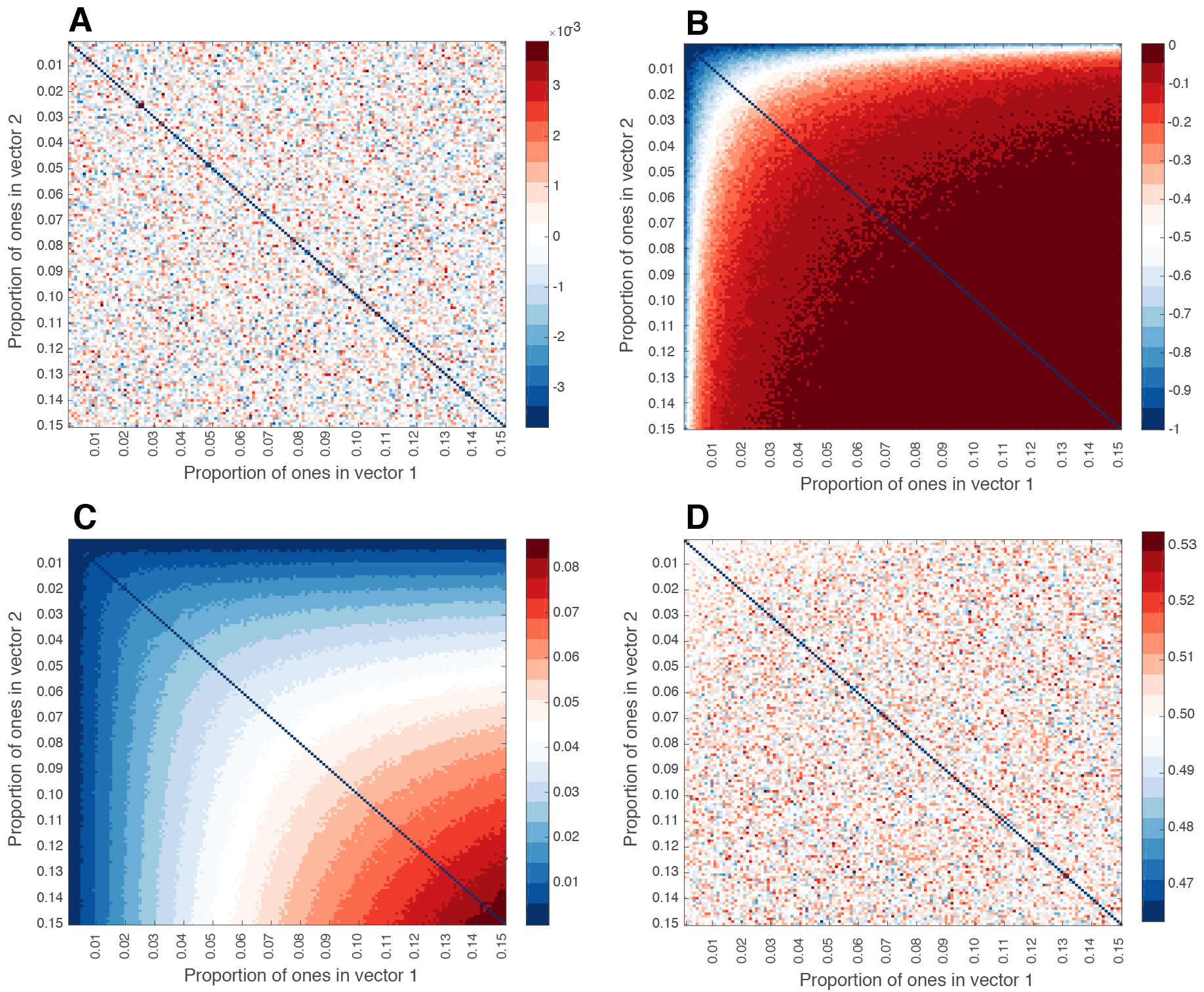
Dependence of correlated gene expression measures on the proportion of positive annotations. We plot the mean value of each metric across 1000 different pairs of random, binary vectors of length 948, which vary only in their proportion of ‘1’s (between 0–0.15; corresponding to a number of ‘1’s ranging from 1 to 150). This is repeated for: (A) mean square contingency coefficient, *r*_*ϕ*_, (B) Jaccard index, (C) Yule’s *Q*, and (D) our developed positive match measure, *p*_match_, Eq. (5). Any systematic trend in correlation values indicates a bias driven by the proportion of positive annotations for a pair of vectors, as is seen for the Jaccard index and Yule’s *Q*. By contrast, *r*_*ϕ*_, which is used through this work, and our probability-based measure, *p*_match_, used to motivate individual gene scoring for enrichment analysis, show no evidence of systematic bias (note the color axis scales).

**Fig S2.**
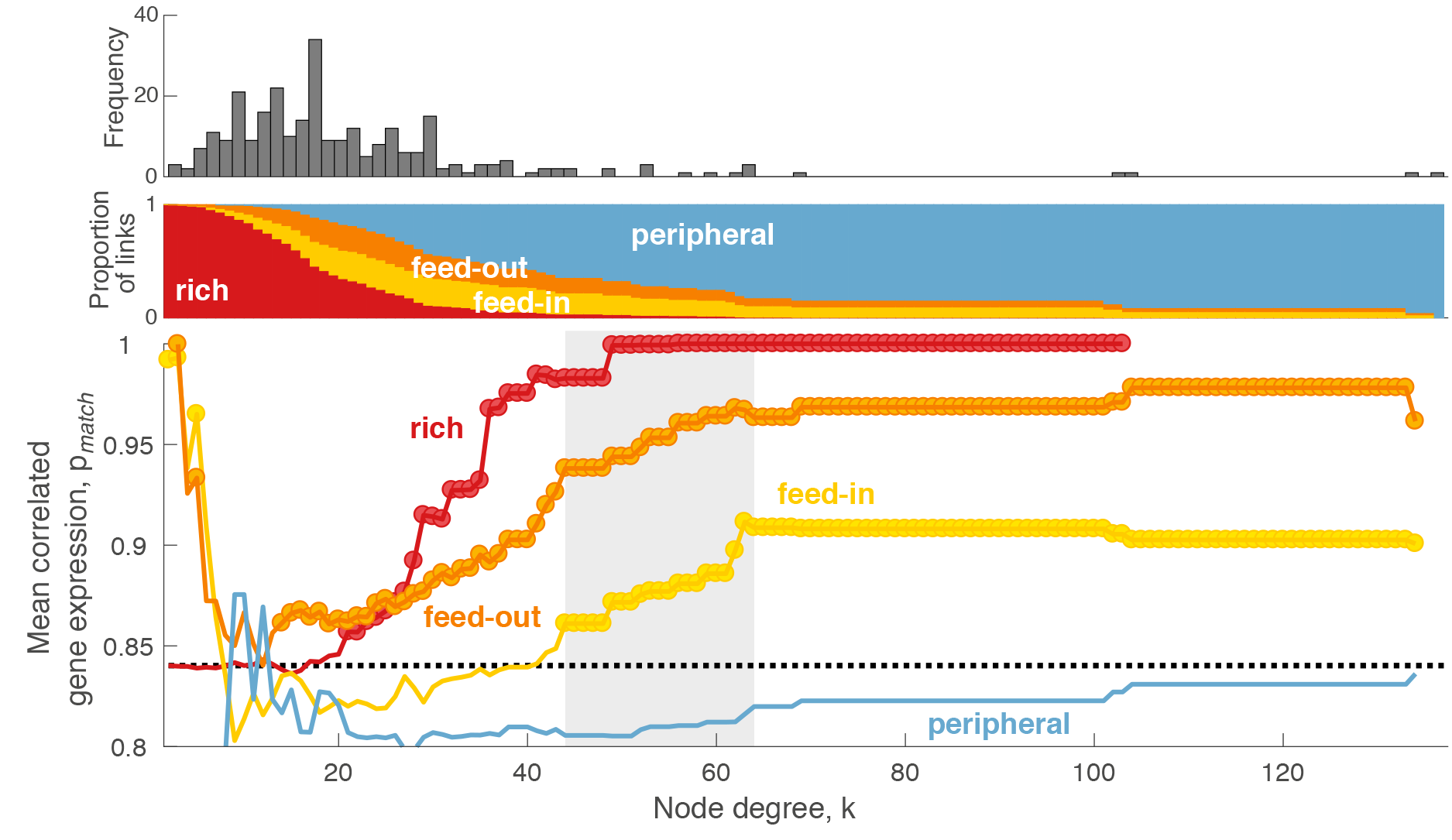
Correlated gene expression measured using the positive matching probability index. The matching probability index, *p*_match_, is introduced here as Eq. (5). *Top*: Degree distribution. *Middle*: Proportion of connections that are ‘rich’ (hub→hub, red), ‘feed-in’ (nonhub→hub, yellow), ‘feed-out’ (hub→nonhub, orange), and ‘peripheral’ (nonhub → nonhub, blue) as a function of the degree threshold, *k*, used to define hubs. Note that at high *k*, most neurons are labeled as nonhubs, and hence the vast majority of connections are ‘peripheral’. *Bottom*: Mean CGE calculated using similarity index from only positive matches, *p_match_*, for each connection type as a function of *k*. The mean CGE across all network links shown as a dotted black line; the topological rich-club regime (determined from the network topology, cf. Fig. 5) is shaded gray. Circles indicate a statistically significant increase in CGE in a given link type relative to the rest of the network (one-sided Welch’s *t* test; *p* < 0.05).

**Fig S3.**
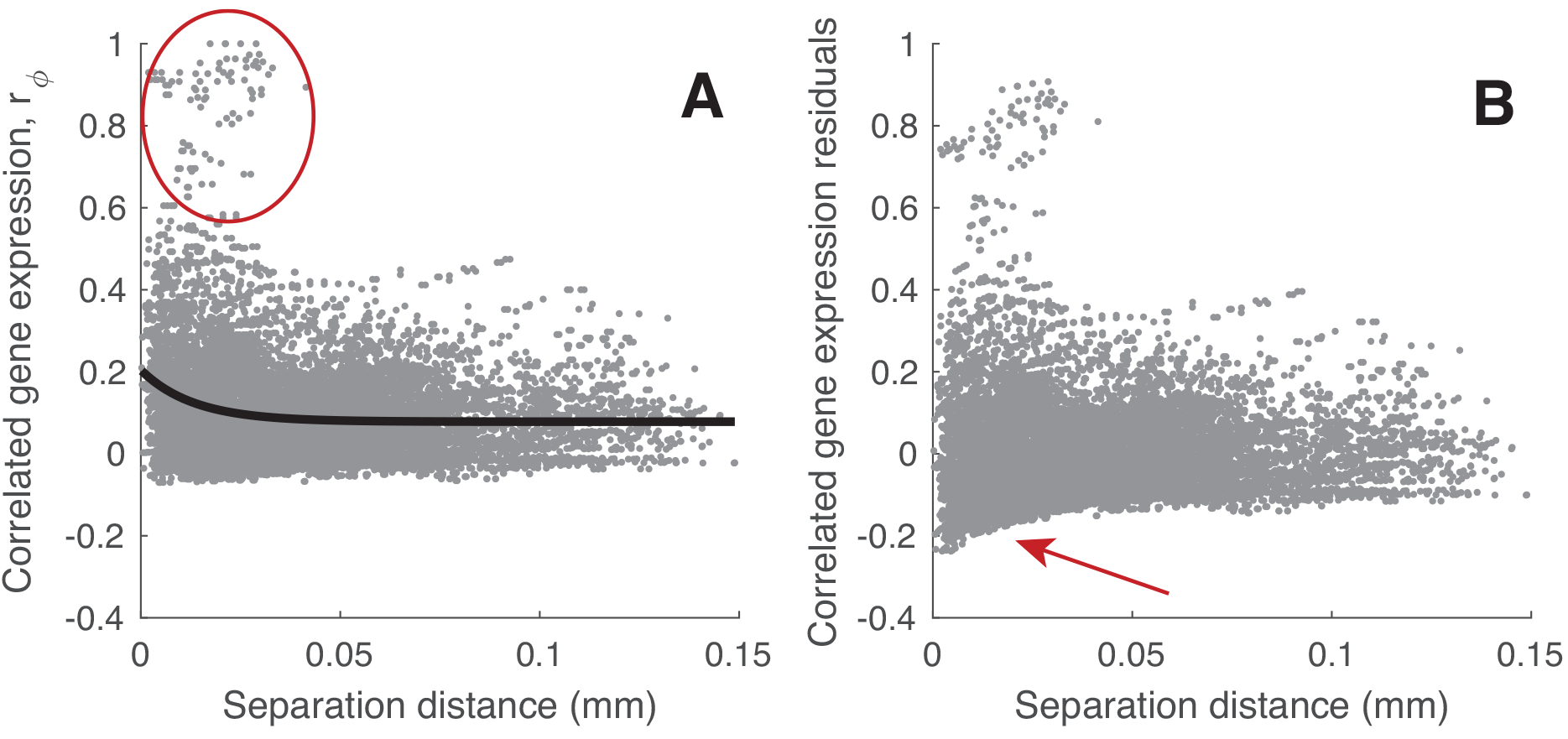
Correcting for spatial effects in CGE data using a bulk exponential trend. Here we consider correlated gene expression in the head, where the strongest spatial relationship exists (cf. Fig. 4. (A) CGE values, *r*_*ϕ*_, plotted as a function of Euclidean separation distance for all pairs of neurons within the head (gray dots), with a fitted exponential trend shown in black, *f* (*x*) = *A* exp(−*λx*) + *B*. (B) Taking residuals from this trend does not adequately correct the spatial trend. Note the artifactual negative correlations indicated with an arrow. This indicates that the trend is not a bulk, isotropic effect, but may instead be driven primarily by a small number of neuron pairs with high *r*_*ϕ*_ at short distances (⪅ 25*μ*m), indicated with a circle in (A).

**Fig S4.**
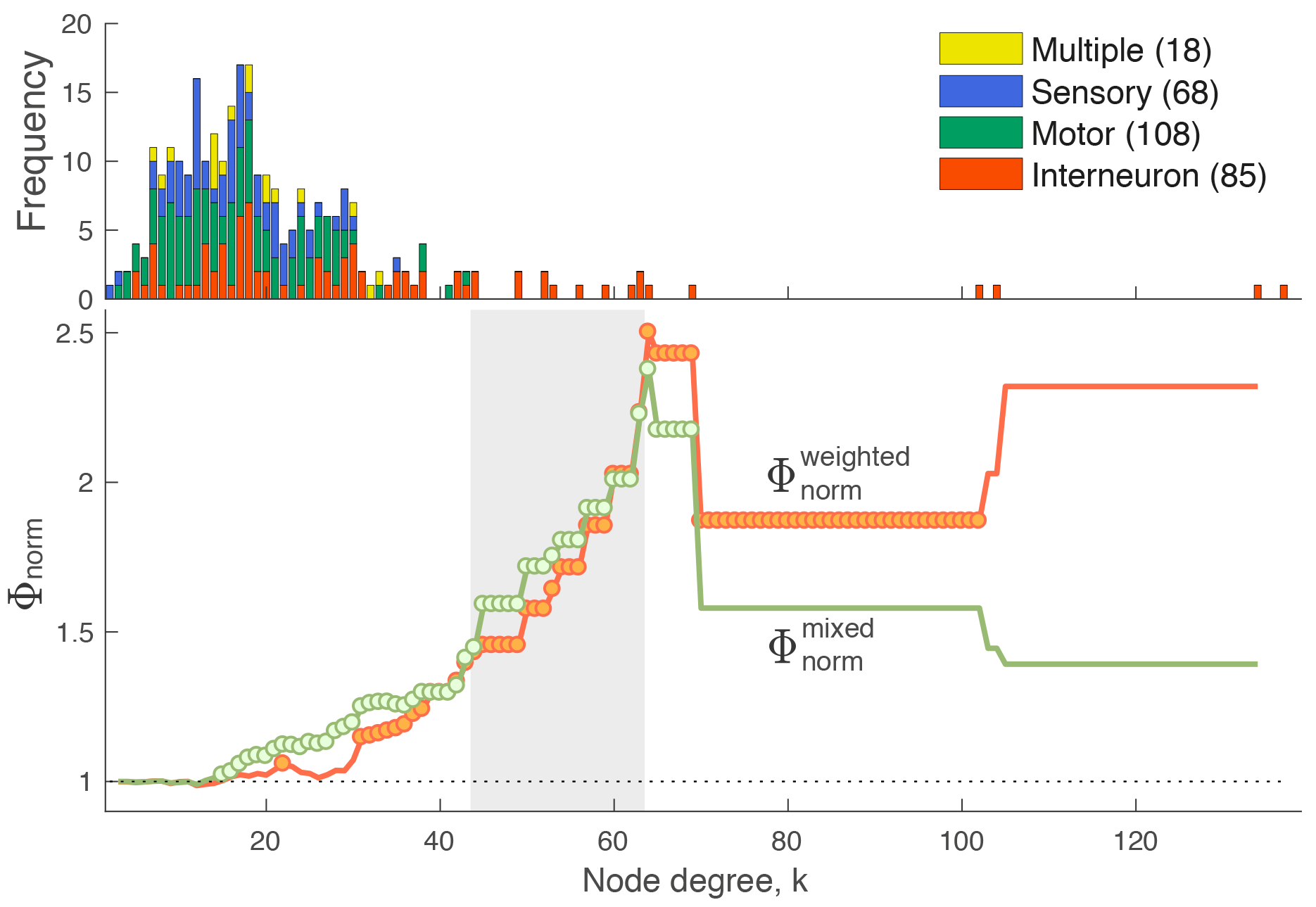
Weighted and unweighted rich-club analyses yield similar results. (A) Degree distribution of the *C. elegans* connectome. Neurons are labeled to four types as in the legend. (B) Normalized weighted rich-club coefficient, 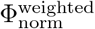 (i.e., topology fixed and weights randomized in the null model, shown orange), and normalized mixed rich-club coefficient, 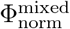 (i.e., both topology and weights mixed in the null model, shown green) are plotted as a function of the degree, *k*, at which hubs are defined (as neurons with degree > *k*) [120]. Circles indicate values of Φ_norm_ that are significantly higher than an ensemble of 1 000 degree-matched null networks (Welch’s *t*-test, *p* < 0.05). Compared to topological rich-club analysis presented in the main text, here the weights of the connections are also accounted for when calculating the rich club coefficient. In the case of the weighted rich-club coefficient, the topology for the null models was kept stable and only the weights of the connections randomized. Results presented here show that connections between higher degree nodes are stronger than expected by chance. On the other hand, in the mixed rich club coefficient both the topology and weights are randomized, therefore we see the combined effect of both types. Distinction between the different null models is discussed in detail in [120]. These results show that connections between high degree nodes are both denser and stronger than expected by chance.

**Fig S5.**
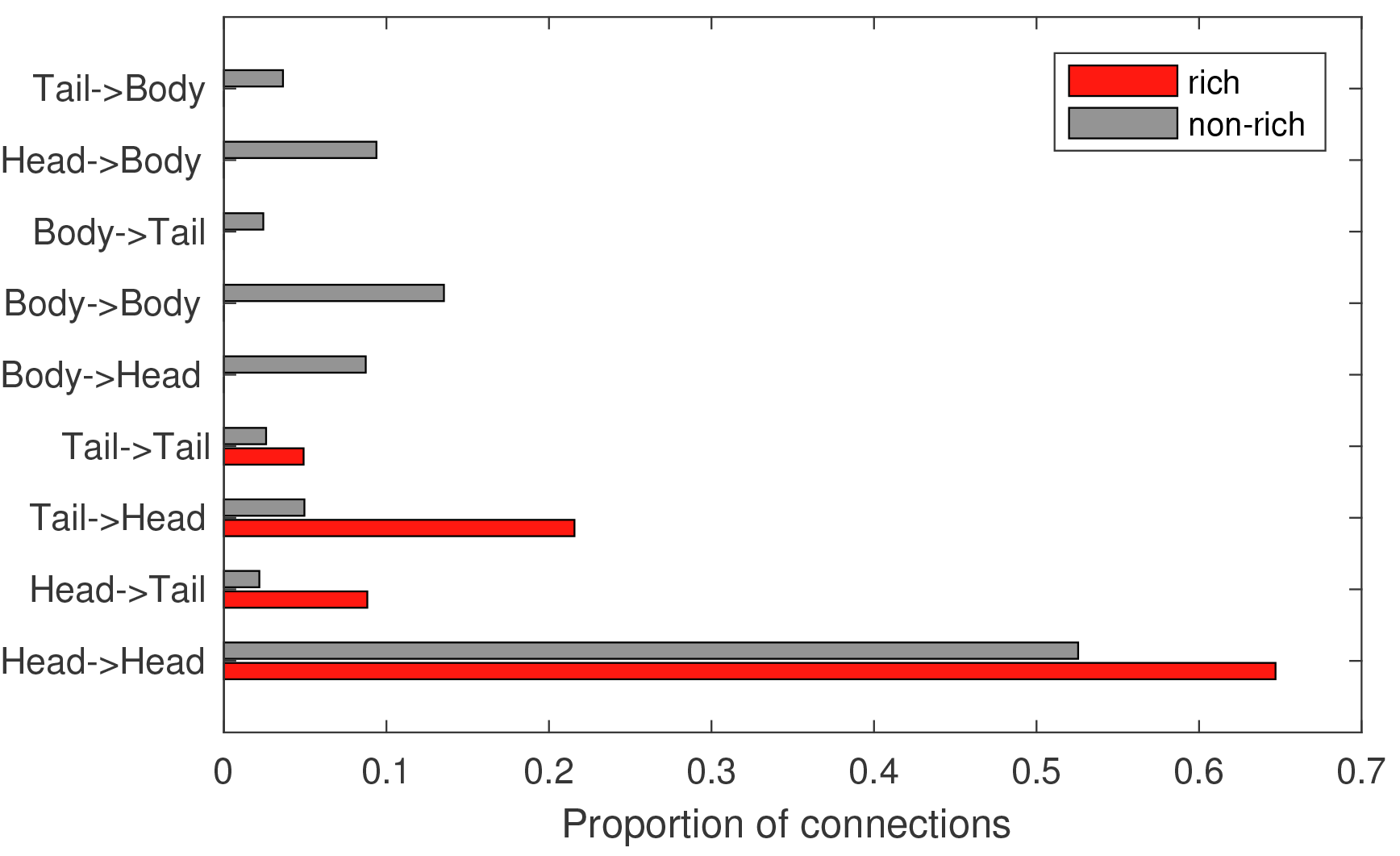
Rich and non-rich connections in the *C. elegans* connectome, categorized by the anatomical location of the source and target neurons. Hub-hub connections (‘rich’) are shown red, and all other connections (‘non-rich’, i.e., feeder and peripheral) are shown gray, where hubs are defined as neurons with degree, *k* > 44. Anatomical locations are labeled as ‘head’, ‘body’, and ‘tail’, and each connection is labeled according to its source and target neurons, labeled on the vertical axis in the form ‘SourceTarget’. The plot shows that the increased separation distance between connected hubs relative to other types of connected neurons is driven by a relative increase in long-range connections between the head and tail.

